# Disrupting the plastid-hosted iron-sulfur cluster biogenesis pathway in *Toxoplasma gondii* has pleiotropic effects irreversibly impacting parasite viability

**DOI:** 10.1101/2022.03.18.484844

**Authors:** Eléa A. Renaud, Sarah Pamukcu, Aude Cerutti, Laurence Berry, Catherine Lemaire-Vieille, Yoshiki Yamaryo-Botté, Cyrille Y. Botté, Sébastien Besteiro

## Abstract

Like many other apicomplexan parasites, *Toxoplasma gondii* contains a plastid harbouring key metabolic pathways, including the SUF pathway that is involved in the biosynthesis of iron-sulfur clusters. These cofactors are key for a variety of proteins involved in important metabolic reactions, potentially including plastidic pathways for the synthesis of isoprenoid and fatty acids. It was shown previously that impairing the NFS2 cysteine desulfurase, involved in the first step of the SUF pathway, leads to an irreversible killing of intracellular parasites. However, the metabolic impact of disrupting the pathway remained unexplored. We have generated another mutant of the pathway, deficient for the SUFC ATPase, and we have investigated in details the phenotypic consequences of TgNFS2 and TgSUFC depletion on parasite homeostasis. Our analysis confirms that Toxoplasma SUF mutants are severely and irreversibly impacted in growth: cell division and membrane homeostasis are particularly affected. Lipidomic analysis suggests a defect in apicoplast-generated fatty acids, along with a simultaneous increase in scavenging of host-derived lipids. However, addition of exogenous lipids did not allow full restauration of growth, suggesting other more important cellular functions were impacted in addition to fatty acid synthesis. For instance, we have shown that the SUF pathway is also key for generating isoprenoid-derived precursors necessary for the proper targeting of GPI-anchored proteins as well as for the parasite gliding motility. Thus, plastid-generated iron-sulfur clusters support the functions of proteins involved in several vital downstream cellular pathways, which implies the SUF machinery may be explored for discovering new potential anti-Toxoplasma targets.

## Introduction

Apicomplexan parasites are some of the most prevalent and morbidity-causing pathogens worldwide. Noticeably, they comprise *Plasmodium* species that can naturally infect humans and cause the deadly malaria in tropical and subtropical areas of the world (1). Although less lethal, another apicomplexan parasite called *Toxoplasma gondii* can cause serious illness in animals, including humans, and has a widespread host range and geographical distribution (2). These protists are obligate intracellular parasites that rely to a large extent on their host cells for nutrient acquisition and for protection from the immune system. Through their evolutionary history, *Plasmodium* and *Toxoplasma* have inherited a plastid from a secondary endosymbiosis event involving the engulfment of a red alga whose photosynthetic capability previously originated from the acquisition of a cyanobacterium (3). Although the ability to perform photosynthesis has been lost during evolution when the ancestors of Apicomplexa became parasitic (4), the plastid has retained critical metabolic functions. For instance it hosts pathways for the synthesis of heme (together with the mitochondrion), fatty acids (via a prokaryotic FASII pathway), isoprenoid precursors (through the so-called non-mevalonate or 1-deoxy-D-xylulose 5-phosphate -DOXP-pathway), and iron-sulfur (FeS) clusters (5, 6). Because of its origin and its metabolic importance, the apicoplast is particularly attractive to look for potential drug targets (7).

As some of the earliest catalytic cofactors on earth (8), Fe-S clusters are found in all kingdoms of life, associated with proteins involved in a number of key cellular functions like the synthesis of metabolites, the replication and repair of DNA, the biogenesis of ribosomes and the modification of tRNAs (9). The biosynthesis of Fe-S clusters necessitates a complex machinery for assembling ferrous (Fe^2+^) or ferric (Fe^3+^) iron and sulfide (S^2-^) ions, and delivering the resulting Fe-S cluster to target client proteins (10). In eukaryotes Fe-S proteins are present in various subcellular compartments like the cytosol and the nucleus, but also organelles of endosymbiotic origin like mitochondria or plastids, and thus require compartment-specific biogenesis systems. The three main eukaryotic Fe-S synthesis pathways comprise the ISC (iron-sulfur cluster) machinery, hosted by the mitochondrion, the cytosolic Fe-S protein assembly (CIA) machinery, important not only for the generation of cytosolic, but also of nuclear Fe-S proteins, and the SUF (sulfur formation) pathway that is found in plastids (9).

Like in plants and algae, apicoplast-containing apicomplexan parasites seem to express the machinery corresponding to the three eukaryotic pathways. For instance, recent investigations in *T. gondii* have shown that the CIA, ISC and SUF pathways are all essential for parasite fitness (11, 12). From a biochemical point of view, mitochondrial and plastidic Fe-S cluster biosynthesis pathways follow a similar general pattern: cysteine desulfurases produce sulfur from L-cysteine, then scaffold proteins provide a molecular platform allowing assembly of iron and sulfur into a cluster, and finally carrier proteins deliver the cluster to target apoproteins. Importantly, targeting the *T. gondii* mitochondrial ISC pathway through disruption of scaffold protein ISU1 was shown to lead to a reversible growth arrest and to trigger differentiation into a stress-resistant form; while on the other hand, targeting the plastidic SUF pathway by inactivating NFS2 function led to an irreversible lethal phenotype (12). Like for many apicoplast-hosted pathways, enzymes belonging to the SUF machinery are essentially absent from the mammalian host and as such they may be seen as good potential drug targets. This has sparked considerable interest for the SUF pathway in *Plasmodium,* which has been shown to be important for the viability of several developmental stages of the parasite (13–17).

To better understand the contribution of the SUF pathway to *T. gondii* viability, we have generated a conditional mutant for the scaffold protein TgSUFC and conducted a thorough phenotypic characterization of this mutant, together with the TgNFS2 mutant we have previously generated (12). Our results confirm that inactivating the plastid-hosted SUF pathway in *T. gondii* leads to irreversible and marked effects on membrane homeostasis, impacting the division process and parasite viability. We show that these effects are likely due to impairment in the function of several key plastidic Fe-S proteins, which have pleiotropic downstream metabolic consequences for the parasite.

## Results

### A Toxoplasma SUFC homolog in the apicoplast

Searching for homologs of the plant SUF system in the ToxoDB.org database, we have previously shown an overall good conservation for the plastidic Fe-S cluster synthesis pathway (12). Among the candidates for members of the SUF machinery, we have identified a potential *T. gondii* homolog of SUFC, member of a Fe-S cluster scaffold complex comprising SUFC, SUFB and SUFD (Fig. 1A). This complex is also present in prokaryotes, where it was first characterized (18): it was shown that bacterial SufC is an ATP-binding cassette (ABC)-like ATPase component essential for proper Fe-S cluster assembly (19). Alignment of the amino acid sequences of the *T. gondii* SUFC candidate (entry TGGT1_225800 in the ToxoDB.org database (20)) and its *Escherichia coli* counterpart showed a good overall conservation (56% of identity), particularly in the motifs that are characteristic of ABC ATPAses (Fig. 1B). The *T. gondii* protein presents a N-terminal extension when compared with *E. coli* SufC, which may contain a transit peptide for targeting to the apicoplast. Accordingly, it was predicted with high probability to be a plastid-localized protein by the Deeploc 1.0 (http://www.cbs.dtu.dk/services/DeepLoc-1.0/) algorithm, although the exact position of the N-terminal transit peptide sequence could not be defined. Data from global mapping of *T. gondii* proteins subcellular location by hyperLOPIT spatial proteomics (21) also suggested an apicoplast localization for TGGT1_225800. To assess whether this protein is a real functional homolog, we first performed complementation assays of an *E.* coli SufC mutant, for which growth is slowed, especially when limiting iron availability with a specific chelator (22). We could show that expressing the predicted functional domain of TGGT1_225800 restored bacterial growth (Fig. 1C), even in the presence of the iron chelator, confirming this protein (hereafter named TgSUFC) is functional.

**Figure 1.**
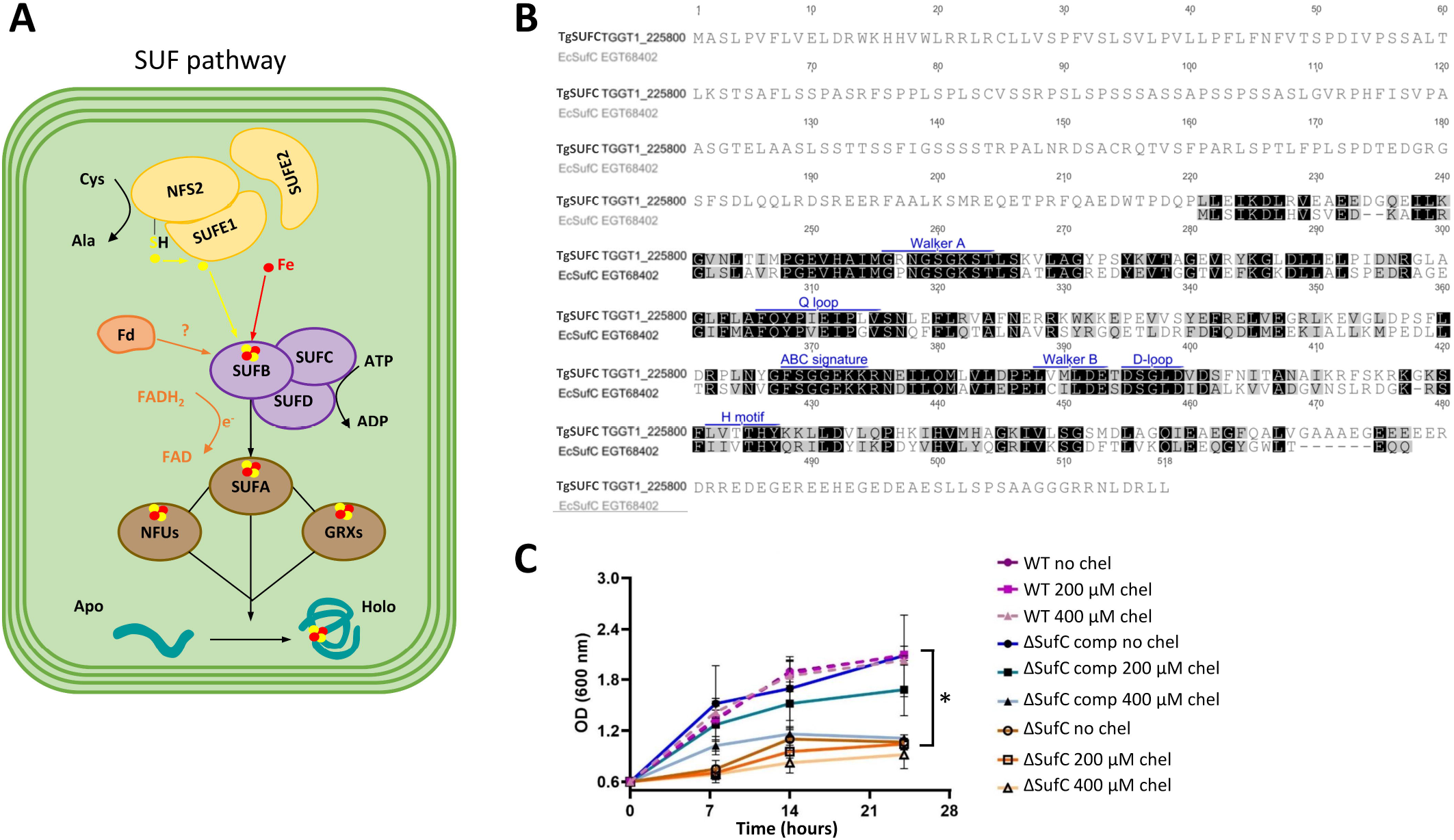
The *Toxoplasma gondii* functional homolog of SufC. A) Schematic representation of the molecular machinery for Fe-S cluster synthesis in the apicoplast of *T. gondii.* B) Alignment of the predicted amino acid sequence of TgSUFC and its homolog from *Escherischia coli.* Motifs that are potentially important for ATPase activity are outlined in blue. C) Functional complementation of bacterial mutants for SufC. Growth of ‘wild-type’ (WT) *E. coli* K12 parental strain, the SufC mutant strain and the mutant strain complemented (‘comp’) by the *T. gondii* homolog, was assessed by monitoring the optical density at 600 nm in the presence or not of an iron chelator (2,2’-bipyridyl, ‘che?). Values are mean from *n* = 3 independent experiments ±SEM. * denotes *p ≤* 0.05, Student’s *t-* test, when comparing values obtained in the absence of chelator for the mutant cell line versus the complemented one.

In order to detect TgSUFC expression and assess its sub-cellular localization in the tachyzoite stage (the fast-replicating stage associated with acute toxoplasmosis (2)), we epitope-tagged the native protein. This was performed by inserting a sequence coding for a C-terminal triple hemagglutinin (HA) epitope tag at the endogenous *TgSUFC* locus by homologous recombination (Figure S1). It was achieved in the TATi ΔKu80 cell line, favoring homologous recombination and allowing transactivation of a Tet operator-modified promoter that we subsequently used for generating a conditional mutant in this background (23–25). Immunoblot analysis with an anti-HA antibody revealed two products, likely corresponding to the immature and mature forms (resulting from the cleavage of the transit peptide upon import into the organelle) of TgSUFC (Fig. 2A). Immunofluorescence assay (IFA) with the anti-HA antibody and co-staining with an apicoplast marker confirmed that TgSUFC localizes to this organelle in *T. gondii* tachyzoites (Fig. 2B).

**Figure 2.**
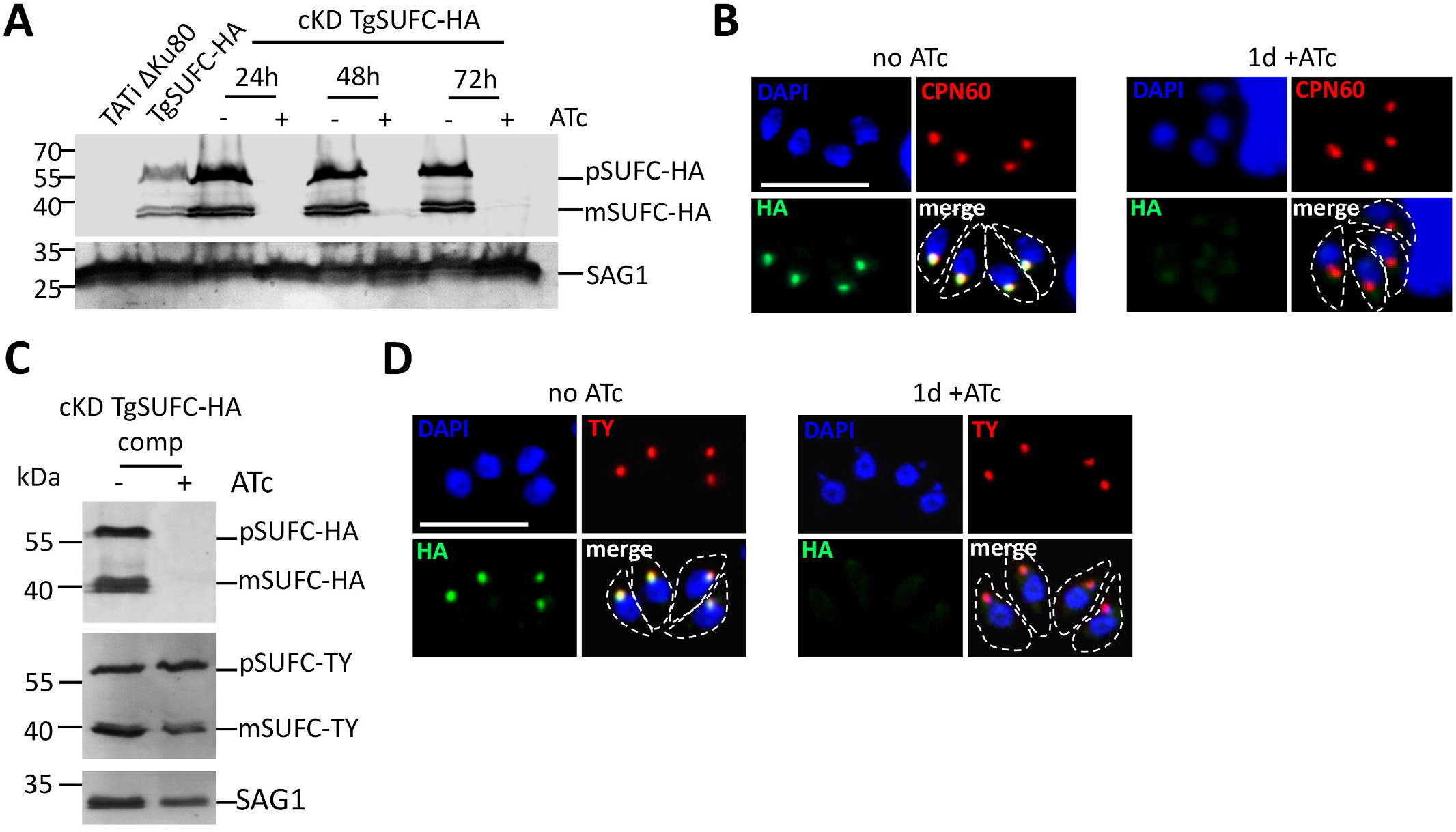
Generation of conditional knock-down and complemented cell lines for the apicoplast-localized TgSUFC. A) Immunoblot analysis with anti-HA antibody shows precursor (p) and mature (m) forms of C-terminally HA-tagged TgSUFC and efficient down-regulation of the protein after 24 hours of incubation with ATc. Anti-SAG1 antibody was used as a loading control. B) HA-tagged TgSUFC (green) localizes to the apicoplast (labelled with marker TgCPN60, red) and is efficiently down-regulated upon addition of ATc for 24 hours. Scale bar represents 5 μm. DNA was labelled with DAPI. Parasite shape is outlined. C) Immunoblot analysis of the conditional TgSUFC knock-down cell line expressing a TY-tagged version of the protein shows similar processing profile and stable expression after 48 hours of ATc addition. D) Immunofluorescence assay confirms co-localization of the regulatable HA-tagged TgSUFC (green) and the TY-tagged additional copy (red), whose expression is retained after 24 hours of incubation with ATc. Scale bar represents 5 μm. DNA was labelled with DAPI. Parasite shape is outlined.

### Depletion of TgSUFC blocks parasite growth

Next, we generated a conditional TgSUFC mutant cell line in the TgSUFC-HA-expressing TATi ΔKu80 background. Replacement of the endogenous promoter by an inducible-Tet07SAG4 promoter was achieved through a single homologous recombination at the locus of interest, yielding the cKD TgSUFC-HA cell line (Fig. S2). In this cell line, the addition of anhydrotetracycline (ATc) can repress transcription through a Tet-Off system (26). Initial phenotypic characterization was performed on two independent clones, which were found to behave similarly and thus only one was analysed further. It should be noted that the promoter replacement resulted in a slightly higher expression of TgSUFC, but did not change the maturation profile of the protein (Fig. 2A). Down-regulation of TgSUFC was assessed by growing the parasites in the presence of ATc. Immunoblot and IFA analyses showed a decrease of TgSUFC to almost undetectable levels after as early as one day of ATc treatment (Fig. 2A and B). We also generated a complemented cell line constitutively expressing an additional TY-tagged (27) copy of TgSUFC from the *uracil phosphoribosyltransferase (UPRT)* locus, driven by a *tubulin* promoter (Fig. S3). This cell line, named cKD TgSUFC-HA comp, was found by immunoblot (Fig. 2C) and IFA (Fig. 2D), to be stably expressing TgSUFC while the HA-tagged copy was down-regulated in the presence of ATc.

We first evaluated the consequences of TgSUFC depletion on parasite fitness *in vitro* by performing a plaque assay, which determines the capacity of the mutant and complemented parasites to produce lysis plaques on a host cells monolayer in absence or continuous presence of ATc for 7 days (Fig. 3). Depletion of TgSUFC prevented plaque formation, which was restored in the complemented cell lines (Fig. 3A and B). Our previous analysis of another SUF pathway mutant (TgNFS2, (12)) suggested that the impact on the pathway leads to irreversible death of the parasites, so we sought to verify this by removing the ATc after 7 days of incubation and monitoring plaque formation. We confirmed that depleting TgSUFC was irreversibly impacting parasite viability, as ATc removal did not lead to the appearance of plaques (Fig. 3C). We next assessed whether this defect in the lytic cycle is due to a replication problem. Mutant and control cell lines were preincubated in absence or presence of ATc for 48 hours and released mechanically, before infecting new host cells and were then grown for an additional 24 hours in ATc prior to parasite counting. We noted that the incubation with ATc led to an accumulation of vacuoles with fewer TgSUFC mutant parasites, but that it was not the case in the complemented cell lines (Fig. 3D). Overall, our data show that depleting TgSUFC leads to an irreversible impact on parasite growth, as previously described for other SUF mutant TgNFS2 (12).

**Figure 3.**
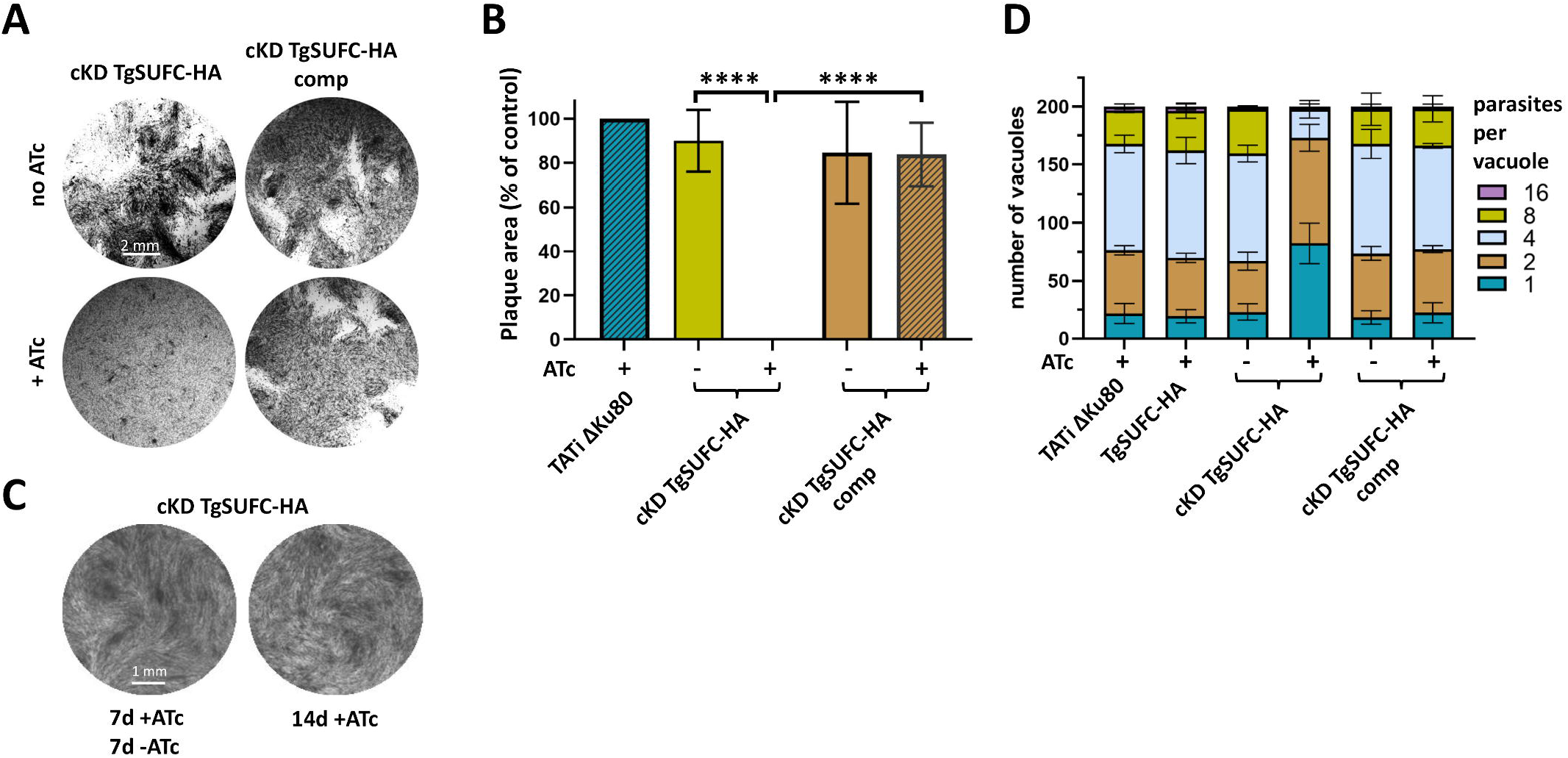
Depletion of TgSUFC affects in vitro growth of the tachyzoites irresversibly. A) Plaque assays were carried out by infecting HFF monolayers with the TgSUFC2-HA conditional knock-down and complemented cell lines. They were grown for 7 days ± ATc. B) Measurements of lysis plaque areas highlight a significant defect in the lytic cycle when TgSUFC is depleted. Values are means of *n =* 3 experiments ± SEM. Mean value of a TATi ΔKu80 control grown in the presence of Atc (not shown on the left) was set to 100% as a reference. **** denotes *p* ≤ 0.0001, Student’s t-test. Scale bar = 2mm. C) Plaque assays for the TgSUFC mutant was performed as described in A), but ATc was washed out after 7 days (7d+ATc 7d-ATc) or not (14d+ATc), and parasites were left to grow for an extra 7 days. No plaque was observed upon Atc removal. Shown are images from one representative out of three independent experiments. Scale bar = 1mm. D) TgSUFC mutant and complemented cell lines, as well as their parental cell line and the TATi ΔKu80 control, were grown in HFF in the presence or absence of ATc for 48 hours, and subsequently allowed to invade and grow in new HFF cells for an extra 24 hours in the presence of ATc. Parasites per vacuole were then counted. Values are means ± SEM from *n* = 3 independent experiments for which 200 vacuoles were counted for each condition.

### SUF pathway mutants display important membrane defects during cell division

*T. gondii* tachyzoites divide by a process called endodyogeny, whereby two daughter cells will assemble inside a mother cell (28). Among the structures which are essential as a scaffold for daughter cell formation is the inner membrane complex (IMC), a system of flattened vesicles underlying the plasma membrane and is supported by a cytoskeletal network. The IMC also supports anchorage for the glideosome, the protein complex powering parasite motility (29). As for several other cellular structures, there is a combination of *de novo* assembly and recycling of maternal material during IMC formation in daughter cells (30). To get more precise insights into the impact of the impairment of the SUF pathway on parasite division, we incubated the TgNFS2 and TgSUFC mutant parasites with ATc for up to 2 days and stained them for IMC protein IMC3 to detect growing daughter cells (Fig. 4A). IMC3 is an early marker of daughter cell budding (31), which is usually synchronized within the same vacuole. However, after two days of ATc treatment an increasing portion of the vacuoles showed a lack of synchronicity for daughter cell budding for both mutant cell lines, although the effect was more pronounced for the TgSUFC mutant (Fig. 4A and B).

**Figure 4.**
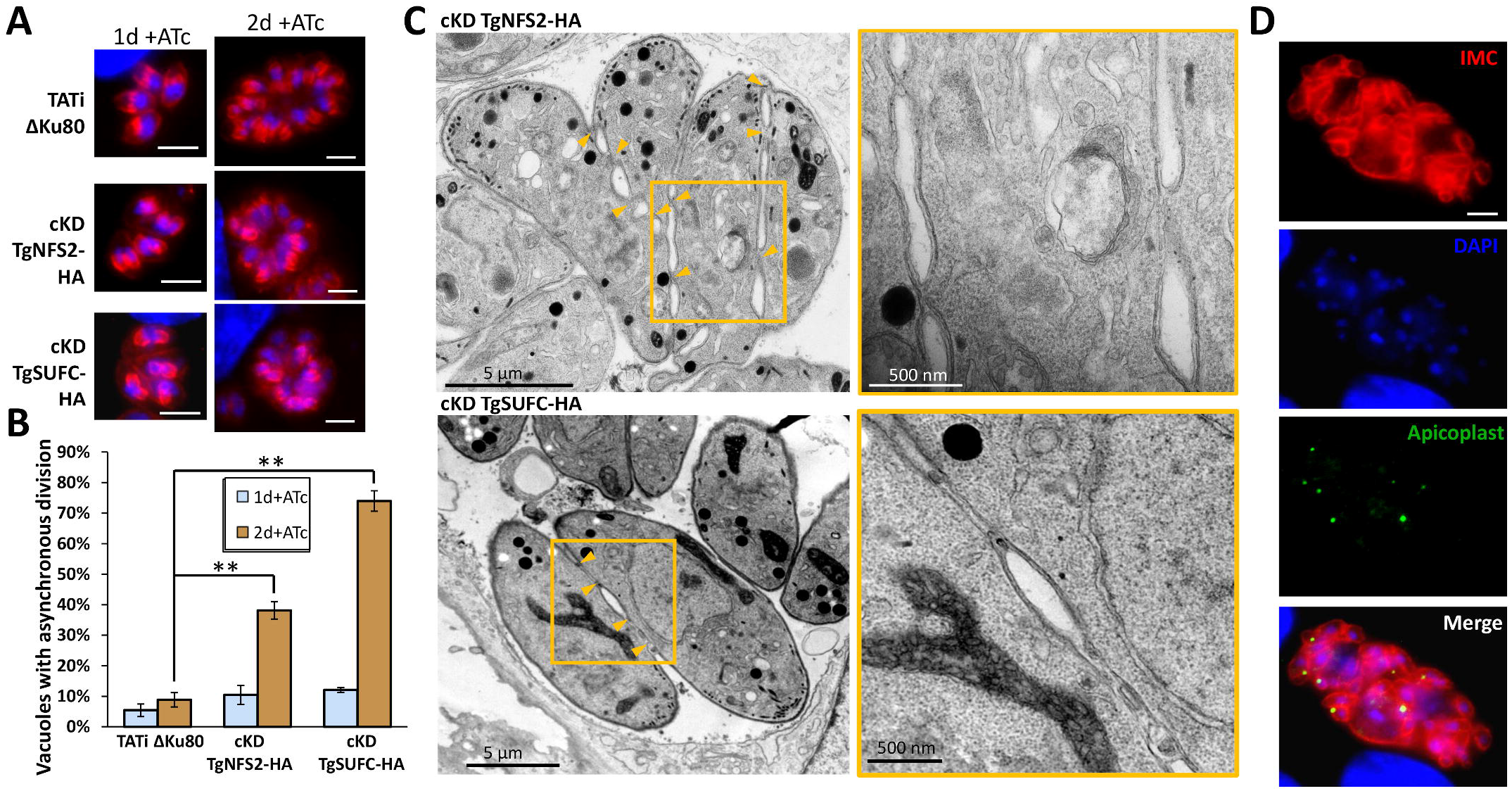
Depletion of TgNFS2 or TgSUFC leads to membrane defects during cell division. A) TgNFS2-HA and TgSUFC-HA conditional knock-down parasites as well as a TATi ΔKu80 control were grown in the presence of ATc for up to 2 days and were stained with an anti-TgIMC3 antibody (in red, to outline parasites and internal buds – top). Scale bar represents 5 μm. DNA was labelled with DAPI (blue). B) The percentage of vacuoles presenting asynchronous division described in B) has been quantified and is represented as a means of *n* = 3 experiments ± SEM (bottom). ** denotes *p* ≤ 0.01, Student’s t-test. C) Electron microscopy analysis of TgNFS2-HA and TgSUFC-HA conditional mutants grown for 4 days in the presence of ATc shows default in plasma membrane separation during parasite division, as displayed on insets representing magnifications of selected parts of the respective left image. D) cKD TgSUFC-HA parasites that were grown in the presence of ATc for 5 days were co-stained with anti-TgIMC3 to outline the inner membrane complex and anti-TgCPN60 (an apicoplast marker), which highlighted abnormal membrane structures and organelle segregation problems. Scale bar represents 5 μm. DNA was labelled with DAPI.

Then, we used electron microscopy to get a subcellular view of the consequences of TgNFS2 and TgSUFC depletion on the cell division process. Strikingly, in parasites grown in the continuous presence of ATc for three days, we observed cytokinesis completion defects. As budding daughter cells emerge, they normally incorporate plasma membrane material that is partly recycled from the mother, leaving only a basal residual body. Here, in both TgNFS2 and TgSUFC mutant cell lines daughter cells remained tethered through patches of plasma membrane (Fig. 4C). Hence, this highlighted an early and important defect in plasma membrane biogenesis and/or recycling during daughter cell budding. We previously observed major cell division defects after long term (five days or more) continuous incubation of cKD TgNFS2-HA parasites with ATc (12). When assessing the cKD TgSUFC-HA parasites in the same conditions, co-staining with apicoplast and IMC markers revealed similar defects, including organelle segregation problems and an abnormal membranous structures (Fig. 4D).

### Depletion of TgSUFC has an impact on the apicoplast

Computational prediction of the Fe-S proteome combined with hyperLOPIT localization data suggests there is a limited number of apicoplast proteins potentially containing Fe-S clusters (12). However, these candidates are supposedly very important for the parasite. They include: IspG and IspH, two oxidoreductases involved in isoprenoid synthesis (32); LipA, a lipoyl synthase important for the function of the pyruvate dehydrogenase (PDH) complex (33); MiaB, which is likely a tRNA modification enzyme (34); as well as of course proteins that are directly involved in Fe-S synthesis, and the plastidic ferredoxin (Fd) that is an important electron donor that regulates several apicoplast-localized pathways (35–37). Dataset from a CRISPR-based genome-wide screen suggests that most of these candidates are important for fitness *in vitro* (38).

We first assessed whether depletion of TgSUFC led to a partial apicoplast loss, as was previously shown for TgNFS2 (12). Although slowed down in growth, some parasites eventually egressed during the course of the experiments and were used to reinvade host cells, and were kept for a total of 5 days in the presence of ATc (Fig. 5A). This is reminiscent to the so-called “delayed death” effect, observed when inhibiting apicoplast metabolism, that often results in slow-kill kinetics (39). Quantification of the apicoplast marker TgCPN60 showed a progressive loss of this protein (Fig. 5A and B). As this could reflect a specific impact on this protein marker rather than a general loss of the organelle, we also stained the parasites with fluorescent streptavidin (mainly detects the biotinylated apicoplast protein acetyl-CoA carboxylase (40)), confirming a similar loss of signal after 5 days of incubation with ATc (Fig. S4). This suggests TgSUFC depletion leads to a progressive but late loss of the apicoplast. Of note, this effect on the organelle is less marked than when TgNFS2 is depleted (12).

**Figure 5.**
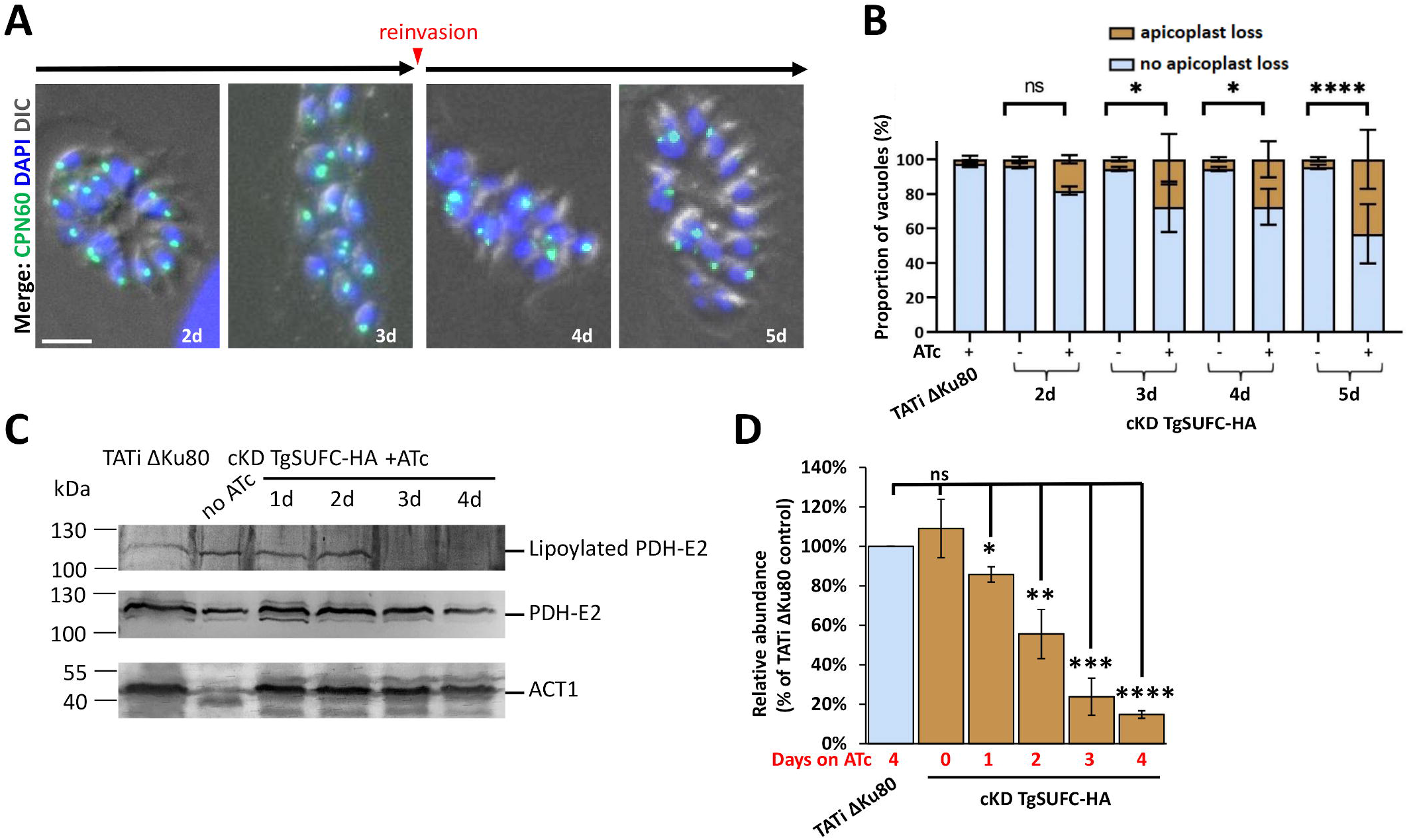
TgSUFC depletion impacts apicoplast-hosted Fe-S pathways. A) cKD TgSUFC-HA parasites were kept in the presence of ATc for up to five days and the aspect of the apicoplast was evaluated by microscopic observation using the specific CPN60 marker. After 3 days, parasites egressed and were used to reinvade new host cells for subsequent timepoints. Scale bar represents 5 μm. DNA was labelled with DAPI. DIC: differential interference contrast. B) Using the labelling described in A), apicoplast loss in vacuoles was monitored after two to five days of incubation with ATc. Data are mean values from *n* = 3 independent experiments ±SEM. ns, not significant; * *p*≤ 0.05, **** *p* ≤ 0.0001, Student’s t-test. C) A decrease in the lipoylation of the E2 subunit of pyruvate dehydrogenase (TgPDH-E2), which depends on the apicoplast-hosted Fe-S-containing lipoyl synthase LipA, was observed by immunoblot using an anti-lipoic acid antibody on cell extracts from cKD TgSUFC-HA parasites kept with ATc for an increasing period of time. A polyclonal antibody raised against PDH-E2 was used as a control for global abundance of the protein and for apicoplast integrity. TgACT1 was used as a loading control. D) Decrease of lipoylated TgPDH-E2 was quantified by band densitometry and normalized with the internal loading control. Data represented are mean ±SEM of *n* = 3 independent experiments, ns, not significant, *≤ 0.05, ** *p*≤ 0.01, *** *p* ≤ 0.001, **** p ≤ 0.0001, ANOVA comparison.

One of the apicoplast-localized Fe-S proteins whose activity can be assessed is LipA, which is responsible for the lipoylation of a single apicoplast target protein, the E2 subunit of the PDH (41). We performed an immunoblot analysis with an anti-lipoic acid on protein extracts from cKD TgSUFC-HA parasites kept in the presence of ATc for up to 4 days (Fig. 5 C and D). We noticed a progressive decrease in lipoylated PDH-E2 to almost no signal after 3 days of ATc incubation. Using an antibody that we specifically raised against the E2 subunit of the PDH, we verified this was not due to a decrease in global levels of this particular protein. Overall, this is comparable to what we previously described upon depletion of TgNFS2 (12). Thus, disrupting the SUF pathway has direct consequences on Fe-S proteins-dependent metabolic pathways hosted by the apicoplast, and prolonged depletion of SUF proteins can even lead to partial loss of the organelle.

### Impact of SUF pathway disruption on fatty acid metabolism

The PDH complex generates acetyl-CoA, which is the first step needed to fuel the FASII system in the apicoplast. This pathway generates fatty acid (FA) precursors that can be subsequently elongated in the endoplasmic reticulum (ER) (42, 43). These de novo synthesized FA from the apicoplast FASII can then be used as essential building blocks, to be combined with scavenged host FA, for bulk phospholipid synthesis to allow essential parasite membrane biogenesis (44, 45). We thus first wanted to evaluate the impact of the perturbation of the SUF pathway on parasite lipid content and homeostasis. Total lipid abundance from the TgNFS2 and TgSUFC mutant cell lines were determined and quantified by gas chromatography-mass spectrometry-based lipidomics analyses (GC-MS). Interestingly, for the TgNFS2 and TgSUFC mutants, there was a significant decrease in the abundance of shorter FAs (C12-C17) (Fig. 6A and D) that was not detected in the complemented strains (Fig. 6B and E). These shorter FA species are usually synthesized via the apicoplast FASII, suggesting that de novo FA synthesis could be affected in these mutants.

**Figure 6.**
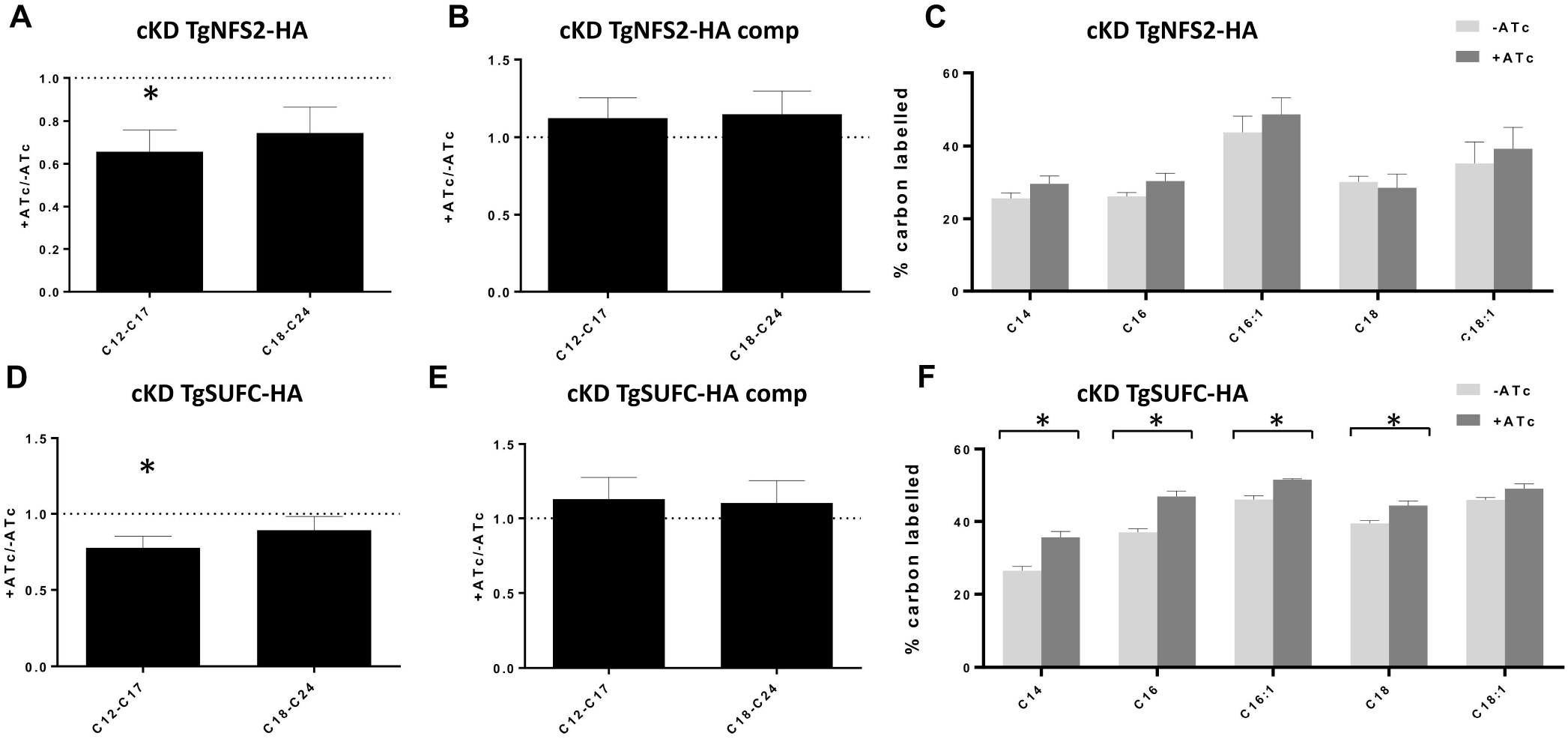
Lipidomic analysis and lipid flux analysis upon TgNFS2 and TgSUFC depletion reveals changes in lipid homeostasis and fluxes. Ratio +ATc/-ATc of total parasite lipid content for cKD TgNFS2-HA mutant (A) and its corresponding complemented cell line (B), and the cKD TgSUFC-HA mutant (D) and its corresponding complemented cell line (E). Host scavenged lipid flux analyses by stable isotope labelling combined to gas chromatography-mass spectrometry analyses on the TgNFS2 (C) and TgSUFC (D) cKD mutants reveal a significant increase of host lipid scavenging upon TgSUFC depletion. All Data are mean from n=4 (or n=3 for panel D) independent experiments ±SEM, * corresponds to a p-value ≤ 0.05 using multiple Student’s t-test.

While FASII was shown previously to be critical for tachyzoite fitness (41), recent investigations have shown that tachyzoites are capable to sense and adapt their lipid synthetic/acquisition capacities according to the host nutrient content and/or lipid availability: for instance they are able upregulate their FASII activity if nutrients are scarce in the host, downregulate it if scavenged lipid levels are too high (45), and scavenge FA precursors from their host cells to at least partly compensate for a lack of *de novo* synthesis (44, 46). Thus, to investigate whether the SUF mutants have their lipid synthesis/flux affected, we sought to assess the impact of TgNFS2 and TgSUFC depletion on *de novo-* synthesized versus scavenged lipids by stable isotope precursor labeling with ^13^C glucose combined with mass spectrometry-based lipidomic analyses (43–45). The analyses revealed a significant increase in the levels of host-scavenged lipids upon the disruption of TgSUFC (Fig. 6F). This was not observed in the complemented cell line (Fig. S5), and thus most likely reflects a mechanism for compensating the lack of de novo-made FAs by increasing scavenging host-derived FAs. On the other hand, this was not as obvious for the TgNFS2 mutant (Fig. 6C).

The ability of tachyzoites to survive lack of de novo lipid synthesis is highly dependent on the availability of exogenous lipid precursors. Others have shown that FASII mutants could be rescued by the addition of palmitic (C16:0) or myristic (C14:0) acid for instance (42, 46, 47). We thus tried to compensate the impact of TgNFS2 and TgSUFC depletion on the parasite lipid homeostasis by supplementing the growth medium with these FAs and performing plaque assays. Palmitic acid supplementation partially restored growth of the TgNFS2 mutant (Fig. 7A and B), while with myristic acid only after long term incubation (up to two weeks) these mutant parasites started growing back (Fig. 7C and D). In contrast, depletion of TgSUFC was not efficiently compensated by exogenous fatty supplementation (Fig. 7). Based on our lipid flux analysis, depletion of TgNFS2 affects the levels of short de novo FA content but mutant parasites are not capable to scavenge more FA from the host membrane lipids (Fig. 6A and C), it is thus possible that providing excess of free FA may help to compensate for this lack of FA and eventually improve fitness of the mutant parasites. On the other hand, TgSUFC mutant parasites do scavenge significantly more FA from the host lipids already (Fig. 6F), and medium supplementation with more exogenous lipids does not seem to further improve their fitness.

**Figure 7.**
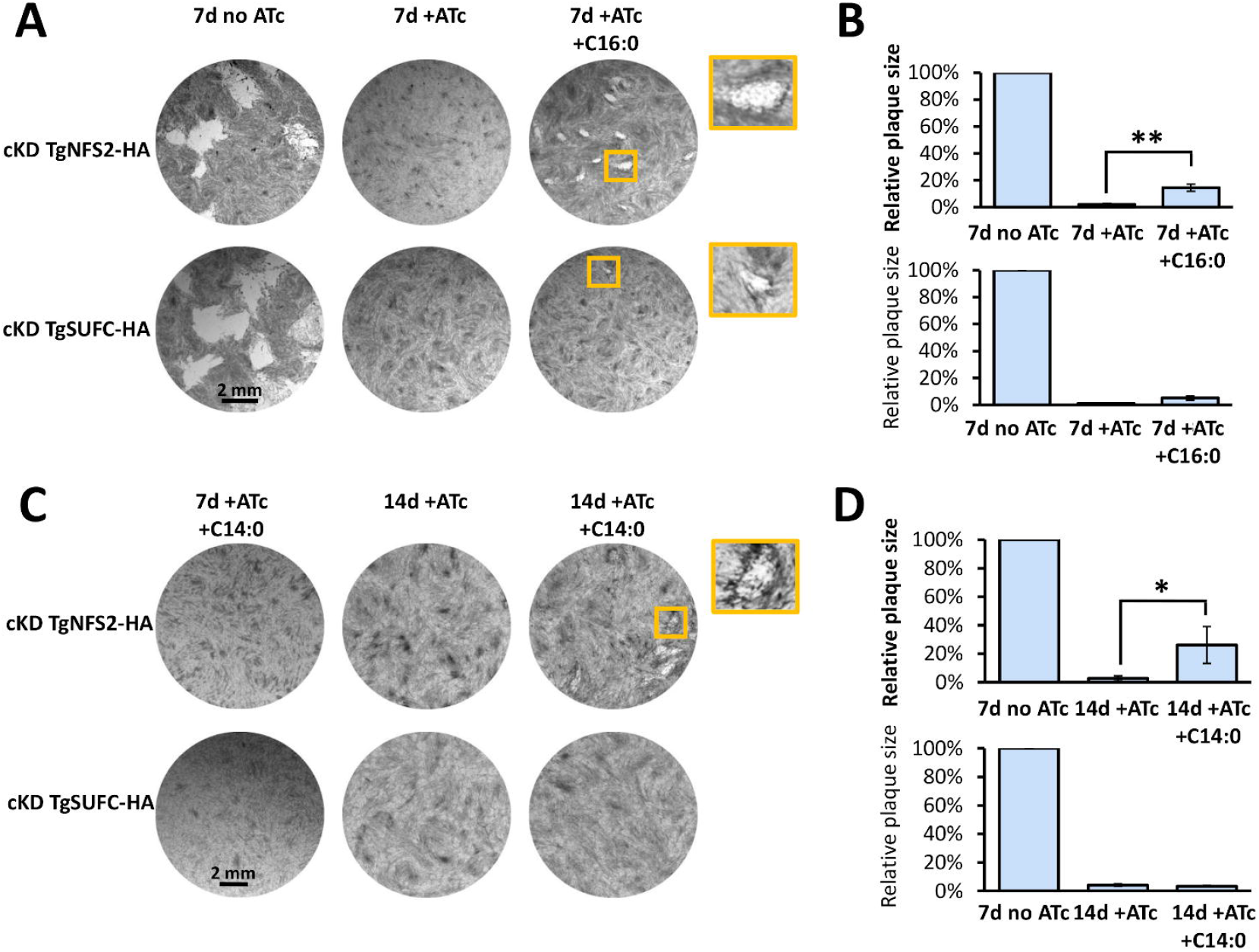
Exogenous supplementation with short chain fatty acids only partially restore fitness of cKD TgNFS2 and cKD TgSUFC mutant parasites *in vitro.* Plaque assays were performed as described in Figure 3A, in the absence or presence of 50 μM of palmitic (C16:0; A, B) or myristic (C14:0; C, D) acid. Plaque sizes were measured and area was expressed as a percentage of the value obtained after 7 days of growth in the absence of ATc. are mean values from n = 3 independent experiments ±SEM. * *p*≤ 0.05, ** *p*≤ 0.01, Student’s t-test. Palmitic acid allows partial restoration of plaques with the cKD TgNFS2 mutant and to a lesser extent with the cKD TgSUFC mutant after 7 days of growth; myristic acid partially restored plaques but only for the cKD TgNFS2 mutant and after two weeks of incubation.

This highlights differences between the two SUF pathway mutants with regards to adaptation to the perturbation of lipid homeostasis, which may be explained by different kinetics in depletion of the respective proteins, or a different impact on global apicoplast function. In any case, our data confirms apicoplast-based lipid production is affected in SUF pathway mutants, and while these parasites establish compensatory mechanisms by scavenging exogenous lipid precursors, they do not allow them recovering to full fitness. This suggests that perturbation of apicoplast FA synthesis is not the only, and likely not even the primary, effect of SUF pathway disruption impacting growth.

### SUF pathway disruption also has an impact on isoprenoid-dependent pathways

The other main apicoplast-localized biosynthetic pathway potentially affected by disruption of the SUF machinery is isoprenoid synthesis, through the two Fe-S-containing proteins IspG and IspH that are needed for the synthesis of the five carbon precursor isopentenyl pyrophosphate (IPP) and its isomer dimethylallyl pyrophosphate (DMAPP) (32). Synthesis of these isoprenoid building blocks is the only essential metabolic function of the apicoplast in the asexual intraerythrocytic stages of *Plasmodium,* where the loss of the organelle can be simply compensated by supplementation with exogenous IPP (48). *Plasmodium* SUF mutants survive when cultured in the presence of IPP, confirming the essential role of the Fe-S synthesizing pathway in this parasite is likely for isoprenoid synthesis (13). Isoprenoid synthesis is also vital for *T. gondii* tachyzoites (49), however IPP supplementation to the culture medium is not possible because, unlike for *Plasmodium,* the highly charged IPP does not efficiently reach the parasite cytoplasm. Tachyzoites can nevertheless scavenge some isoprenoids precursors from their host cells (50). Apicoplast-derived isoprenoid precursors are mostly known for their involvement in important posttranslational protein modifications like prenylation, glycosylphosphatidylinositol (GPI) anchoring, as well glycosylation, in addition to the synthesis of quinones and several antioxidant molecules (32).

In *Plasmodium* it is believed that prenylated proteins that regulate vesicle trafficking are key in the delayed death phenotype caused by apicoplast loss (51). As geranylgeraniol (GGOH), an isoprenoid precursor for protein farnesylation/prenylation, was successfully used to at least partially complement deficiencies in apicoplast isoprenoid production (50, 52), we tried to supplement the culture medium with GGOH and perform plaque assays with the SUF mutants we generated. However, we did not detect any restoration of growth (Fig. S6A). In contrast to *Plasmodium* (53, 54), the prenylome of *T. gondii* is still largely uncharacterized, but using an anti-farnesyl antibody, we did not detect obvious alterations in the general profile of prenylated proteins upon depletion of TgNFS2 or TgSUFC (Fig. S6B). Although apicoplast-generated IPP and DMAPP are also necessary for synthesizing the farnesyl diphosphate used for protein prenylation in *T. gondii* tachyzoites, it is thus possible that these parasites can initially scavenge host-derived isoprenoids to compensate for a deficient *de novo* production (50). In any case, altogether our results suggest that defects in protein farnesylation/prenylation may not be one of the primary consequences of SUF pathway disruption in *T. gondii.*

We next sought to investigate the potential impact of TgNFS2 or TgSUFC depletion on GPI anchoring. The SAG-related sequence (SRS) family comprising proteins related to SAG1, the first characterized *T. gondii* surface antigen, is arguably the best characterized family of GPI-anchored proteins in the parasite (55). We pre-incubated mutant parasites for three days with ATc, and allowed them to invade and grow into host cells for an extra day in the presence of ATc before using specific antibodies to detect GPI-anchored SAG1 and SAG3. We could see obvious signs of mislocalization for these two proteins that, instead of keeping a homogenous peripheral distribution, they were often seen accumulated at the apex or base of the parasites, or found within the parasitophorous vacuole space (Fig. 8A and B). Interestingly, while the SAG1 protein appears to be distributed differently by IFA, our previous immunoblot analyses suggest there is no drastic change in the total amount of protein upon TgNFS2 or TgSUFC depletion ((12) and Fig. 2A). It was previously shown that the deletion of SAG1’s GPI anchor leads to a constitutive secretion of this protein to the parasitophorous vacuole space (56). It should be noted that, in contrast to the distribution of GPI-anchored SAG proteins, in these experimental conditions the overall structure of the IMC appeared unaffected (Fig. 8A and B). Hence, our data suggest that disruption of the SUF pathway perturbs GPI anchor formation.

**Figure 8.**
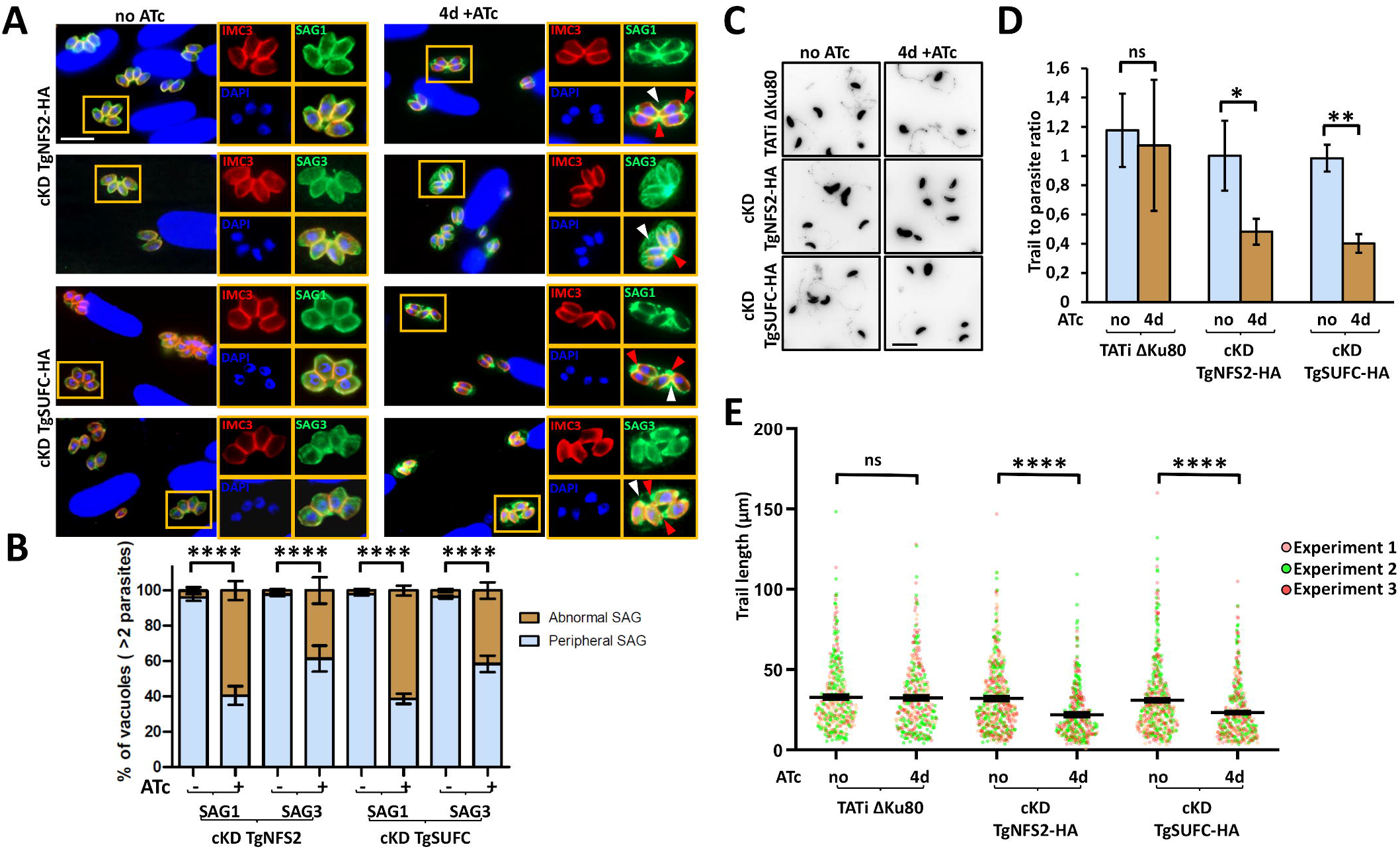
The depletion of TgNFS2 or TgSUFC leads to mislocalization of GPI-anchored surface antigens and impacts gliding motility. A) TgNFS2 and TgSUFC conditional mutants were grown for three days in the presence or absence of ATc and allowed to invade host cells for another 24 hours in the presence or absence of ATc. Parasites were then co-stained for inner membrane complex marker IMC3 (red) together with GPI-anchored protein SAG1 or SAG3 (green). As shown on insets representing selected parts of the images, the depletion of TgNFS2 or TgSUFC leads to the accumulation of SAGs in the vacuolar space (white arrowhead) or concentration at the apex or base of the parasite (red arrowhead). Scale bar represents 10 μm. DNA was labelled with DAPI. B) Quantification of the abnormal distribution of SAG labellings in vacuoles containing more than two parasites. Data are mean values from *n* = 3 independent experiments ±SEM. **** *p* ≤ 0.0001, Student’s t-test. C) Representative views of a gliding assay showing lower abundance of SAG1 trails upon TgNFS2 or TgSUFC depletion (inverted grayscale images). Scale bar represents 10 μm. D) Quantification of the trail to parasite ratio on at least ten randomly selected fields. Data are mean values from *n* = 3 independent experiments ±SEM. ns, not significant; * *p*≤ 0.05, ** *p* ≤ 0.01, Student’s t-test. E) Individual measurements of SAG1 trail lengths. Horizontal lines represent mean values from *n* = 3 independent experiments ±SEM. ns, not significant; **** *p* ≤ 0.0001, Student’s *t*-test.

Another important posttranslational modification depending on isoprenoid-containing dolichol is glysosylation. Several key proteins of the glideosome complex are supposedly glycosylated, and as a consequence glycosylation inhibition has been reported to impact parasite motility (57, 58). We thus performed gliding motility assays on the SUF mutants. Typically, this is done by monitoring the shedding of SAG1, that leaves trails when tachyzoites glide on solid substrates. Perhaps as a consequence of SUF pathway disruption on SAG1 targeting, trails they were clearly less abundant upon depletion of TgNFS2 and TgSUFC (Fig. 8C and D). We could nevertheless detect and measure trails, whose mean length provides an estimate of overall motility rates, and they were found to be significantly smaller in absence of TgNFS2 or TgSUFC (Fig. 8E). These results suggest that protein glycosylation is also affected upon disruption of the SUF pathway. Overall, our findings indicate that the depletion of proteins of the SUF pathway leads to defects in the synthesis of isoprenoids precursors with consequences on the post-translational modification and targeting of proteins of the peripheral membrane system.

## Discussion

Beside components of the SUF Fe-S cluster synthesis machinery, the apicoplast harbors only a small number of putative Fe-S proteins (Fig. 9, (12)). Yet, they are all presumably important for parasite fitness as suggested by their negative scores with a CRISPR-based genome-wide screen (Fig. 9, (38)). However, not all are expected to be absolutely essential *in vitro.* The tRNA modifying enzyme MiaB for instance, has only a moderately low fitness score and has been shown recently in to be dispensable for *Plasmodium* intraerythrocytic stages (37). Similarly, the LipA lipoyl synthase essential for the function of the E2 subunit of the PDH complex, which is in turn crucial for generating the acetyl-CoA necessary for de novo FA synthesis in the apicoplast, seems dispensable for *Plasmodium* blood stages (37), for which the FASII system is not essential in high nutrient-content medium (44, 59). However, given the potentially greater importance of FASII in *T. gondii* tachyzoites (41), this is something we investigated further. We demonstrated disruption of the SUF pathway impaired LipA function and PDH-E2 lipoylation (Fig. 5) and, likely as a consequence of this and FASII perturbation, general production of myristic and palmitic acid in the parasites (Fig. 6). However, the growth defect of SUF mutants could only be partially complemented by FA supplementation for the TgNFS2 mutant, and not at all for the TgSUFC mutant (Fig. 7), suggesting perturbation of FASII is not the primary cause of death for these mutants. While it is undoubtedly a metabolic pathway that plays a central role for parasite fitness, the view on the essentiality of FASII in *T. gondii* tachyzoites has recently evolved. There is now published evidence that parasites can adapt their metabolic capacities depending on the nutrient environment (44, 45), and even survive *in vitro* when FASII enzymes are inactivated (46, 60). There is clearly flexibility in the adaptation of parasite pathways to lipid sources (44, 45, 61). As tachyzoites can readily scavenge and incorporate FAs from exogenous sources (i.e. phospholipid made by the host cell, and/or phospholipids and FA scavenged from the extracellular medium) into their own range of lipids (44–46, 62), in the end the essentiality of the FASII pathway depends largely on nutrient availability *in vivo,* or *in vitro* through culture conditions provided.

**Figure 9.**
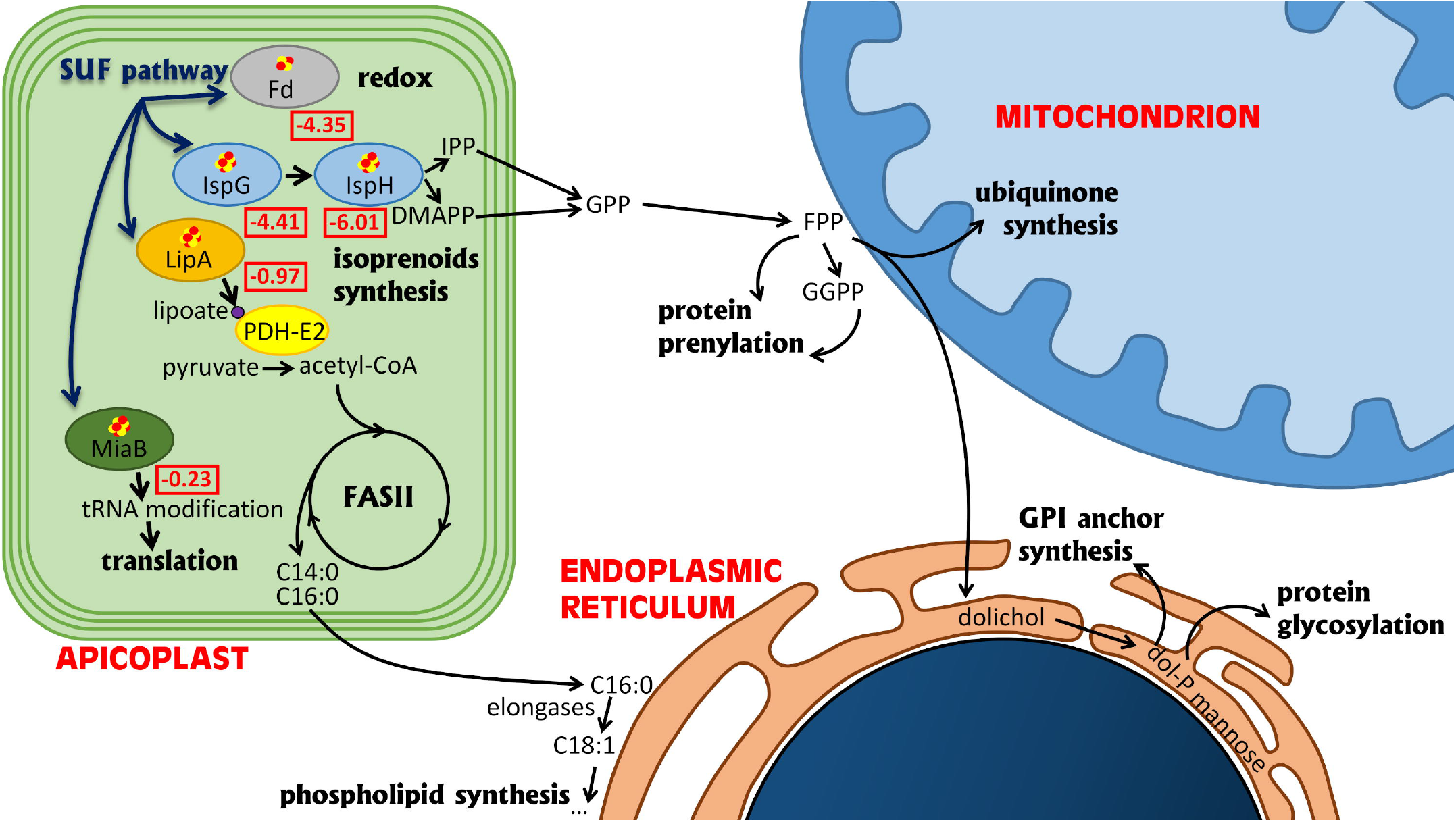
Schematic representation of th main celllular pathways that depend on apicoplast Fe-S proteins. For selected apicoplast-located Fe-S proteins squared red numbers represent CRISPR fitness score of the corresponding gene (genes that contribute to *in vitro* parasite fitness are represented by negative scores; values below −2.5 highlight increasing likelihood of being essential). Fd: ferredoxin; IspG/lspH: oxidoreductases catalysing the last two steps of IPP/DMAPP synthesis; LipA: lipoyl synthase; PDH-E2: E2 subunit of the pyruvate dehydrogenase complex; CoA: coenzyme A; IPP: isopentenyl diphosphate; DMAPP: dimethylallyl diphosphate; FASII: fatty acid synthesis type II; GPP: Geranyl diphosphate; FPP: farnesyl diphosphate; GGPP: Geranylgeranyl diphosphate; GPI: Glycosylphosphatidylinisotol; dol-P: dolichol phosphate.

The other main metabolic pathway that depends directly on Fe-S proteins in the apicoplast is for the synthesis of isoprenoid precursors. When their isoprenoid production is inhibited, *T. gondii* tachyzoites can scavenge some of these precursors from the host cell, leading to a delayed death effect (50, 63), but that cannot fully compensate for a lack of de novo synthesis. Isoprenoids are a large and diverse class of lipids whose cellular functions in Apicomplexa still remain to be extensively characterised, but they include synthesis of vitamins and cofactors (ubiquinone), and are also involved in important posttranslational modifications of proteins, like prenylation, GPI-anchoring and glycosylation (Fig. 9, (32)). Ubiquinone is a central molecule in the mitochondrial electron transport chain (ETC): its quinone head is the functional group for the transfer of electrons, whereas the isoprenoid tail primarily serves for anchoring to the inner mitochondrial membrane. The mitochondrial ETC is a validated drug target in Apicomplexa, for which complex III inhibitor atovaquone has been used in therapeutic strategies (64). However, while *T. gondii* mitochondrial ETC mutants are severely impaired in growth (65–68), it seems that genetic or pharmaceutical inactivation (with atovaquone) is reversible (12, 67), and may lead to stage conversion and a metabolic dormant state rather than complete death of the parasites. On the contrary, the viability of the SUF mutants is irreversibly affected (Fig. 3C and (12)). So although the impact SUF protein depletion on the isoprenoid pathway is likely to lead to a deficiency in ubiquinone synthesis, which in turn would contribute to a decrease in parasite fitness, this is probably not the main reason for the irreversible death of the parasites.

In *Plasmodium* blood stages, where isoprenoid synthesis is the only essential pathway hosted by the apicoplast, disrupting the prenylation of Rab GTPAses, which are involved in vesicular trafficking, contributes to delayed death (51). In *T. gondii,* interestingly, perturbation of Rab function can lead to intracellular accumulation and patchy surface distribution of SAG1 proteins, and results in defects in the delivery of new membrane required for completing daughter cell segregation at the end of cytokinesis (69, 70). This bears some similarity with some phenotypes we have observed in the SUF mutants, yet while we cannot completely exclude Rab prenylation is perturbed in the SUF mutants, we did not identify any particular prenylation problem in the parasites, and we failed to complement their growth defect with prenylation precursor GGOH (Fig. S6). We thus investigated other important isoprenoid-dependent protein modifications. For instance, upon sugar addition dolichol can be used for the formation of GPI anchors, or act as a donor for protein glycosylation. Our phenotypic analysis revealed that the targeting of GPI-anchored surface proteins and the gliding motility of the parasites, which relies on glycosylated proteins, were clearly affected upon disruption of the SUF pathway (Fig. 8). This confirms the importance of the apicoplast Fe-S cluster synthesis machinery for isoprenoid metabolism. In *Plasmodium,* interfering with apicoplast-hosted isoprenoid production affects the morphology of the organelle (71) but depletion of IspG and IspH does not lead to loss of the apicoplast (37), while interfering with the SUF pathway does (13). We also observe a late impact on the organelle (Fig. 5), that suggest the phenotypic consequences of SUF proteins in *T. gondii* are indeed multifactorial and would extend beyond the simple disruption of the isoprenoid pathway.

At the cellular level, one of the most visible consequence of long term depletion of SUF proteins is the membrane defects in the late stages of cytokinesis (Fig. 4). Interestingly, treatment of tachyzoites with the FASII inhibitor triclosan or inactivating the FASII component acyl carrier protein was shown to lead to severe problems in cytokinesis completion, with tethered daughter cells resembling the phenotype we have described here (47). A similar phenotype was also observed in the mutant for TgATS2, an apicoplast-located acyltransferase responsible for phosphatidic acid synthesis (44). This points to a central role for the apicoplast to provide specific precursors for membrane biogenesis during cytokinesis, and to a SUF-dependent FASII function is important for the homeostasis of the parasite plasma membrane. It should however be noted that some isoprenoid-dependent cellular mediators may also contribute to plasma membrane synthesis during cell division. For instance, disruption of Rab-controlled vesicular trafficking, leads to very similar phenotypes of incomplete cytokinesis, with tachyzoites still fused along their lateral surface (69, 70). Glycosylated IMC proteins associated with gliding motility are also important for IMC formation and the cell division process (72). It is also possible that yet unidentified GPI-anchored *T. gondii* proteins may be involved in plasma membrane formation or recycling: GPI synthesis is essential for *T. gondii* survival (73), but the function of individual GPI-anchored proteins remain largely overlooked. The detrimental effects of SUF depletion on plasma membrane homeostasis may thus manifest through both FASII and isoprenoid perturbation. Moreover, as some isoprenoid-dependent modifications are also linked to FA acid synthesis, like GPI anchors of *T. gondii* surface proteins that also necessitate phospholipid moieties (74), a simultaneous impact on the two pathways may enhance the phenotypic output.

One key apicoplast-located Fe-S protein is Fd, which has a central role in the function of Fe-S-dependent apicoplast enzymes: it is potentially providing electrons to other apicoplast Fe-S enzymes like MiaB, IspG, IspH and LipA. The role of Fd has recently been investigated in Apicomplexa. In *Plasmodium* blood stage, the loss of parasite viability upon Fd depletion was likely mostly due to the importance of Fd for the isoprenoid synthesis pathway (37), which is the only essential apicoplast-located pathway in this developmental stage. Fd is equally essential for *T. gondii,* tachyzoite survival, where it was shown to impact both FASII and isoprenoid synthesis (35), in a similar fashion to our SUF mutants. Whether the vital importance of Fe-S cluster synthesis and associated apicoplast redox metabolism is solely through its key role in isoprenoid synthesis is thus less clear in *T. gondii* than in *Plasmodium.* As *T. gondii* tachyzoites grow in host cell types that can potentially provide them with more resources, the ability to scavenge exogenous metabolites creates a complex situation whereby metabolic pathways like FASII may be only essential in certain particular conditions. The nutrient-rich in vitro culture systems may also mask some important contributions. In any case, because of their upstream role in cellular functions important for parasite fitness, Fd and SUF mutants clearly have pleiotropic effects. More importantly, we have confirmed here that disrupting the SUF machinery leads to an irreversible death of the tachyzoites, which is not the case, for example, when the mitochondrial Fe-S cluster machinery is inactivated (12). For all these reasons, and also because of its absence from mammalian hosts of the parasite, the SUF pathway has a strong potential identifying novel drug targets.

## Experimental procedures

### Parasites and cells culture

Tachyzoites of the TATi ΔKu80 *T. gondii* strain (25), as well as derived transgenic parasites generated in this study, were maintained by serial passage in monolayers of human foreskin fibroblast (HFF, American Type Culture Collection, CRL 1634) grown in Dulbecco’s modified Eagle medium (Gibco), supplemented with 5% decomplemented fetal bovine serum, 2-mM L-glutamine and a cocktail of penicillin-streptomycin at 100 μg/ml.

### Bioinformatic analyses

Sequence alignment was performed using the Multiple Sequence Comparison by Log-Expectation (MUSCLE) algorithm of the Geneious 6.1.8 software suite (http://www.geneious.com). Transit peptide and localization prediction was done with the Deeploc 1.0 (http://www.cbs.dtu.dk/services/DeepLoc-1.0/) algorithm.

### Heterologous expression in *E. coli*

Construct for designing the recombinant protein used for *E. coli* complementation was defined by aligning the amino acid sequences of TgSUFC with its *E. coli* counterparts. A 894 bp fragment corresponding to amino acids 220-518, was amplified by polymerase chain reaction (PCR) from *T. gondii* cDNA using primers ML4200/ML4010 (sequences of the primers used in this study are found in Table S1). The fragment was cloned into the pUC19 plasmid (Thermo Fisher Scientific) using the HindIII/BamHI restriction sites. The SufC *E. coli* mutant from the Keio collection (obtained from the The *Coli* Genetic Stock Center at the University of Yale: stain number JW1672-1), was transformed with the plasmid expressing the TgSUFC recombinant protein and selected with ampicillin. For growth assays (22), overnight stationary phase cultures were adjusted to the same starting OD_600_ of 0.6 in salt-supplemented M9 minimal media containing 0.4% glucose and varying amounts of the 2,2□-BipyridyI iron chelator (Sigma-Aldrich). Growth was monitored through OD_600_ measurement after 7,14 and 24 hours at 37°C in a shaking incubator.

### Generation of the HA-tagged TgSUFC cell line

A CRISPR-based strategy was used. Using the pLIC-HA_3_-CAT plasmid as a template, a PCR was performed with the KOD DNA polymerase (Novagen) to amplify the tag and the resistance gene expression cassette with primers ML3980/ML3981, that also carry 30□bp homology with the 3□ end of the corresponding genes. A specific single-guide RNA (sgRNA) was generated to introduce a double-stranded break at the 3□ of the *TgSUFC* gene, using primers ML3952/ML3953, and the protospacer sequences were introduced in the Cas9-expressing pU6-Universal plasmid (Addgene, ref #52694) (38). The TATi ΔKu80 cell line was transfected and transgenic parasites were selected with chloramphenicol and cloned by serial limiting dilution.

### Generation of TgSUFC conditional knock-down and complemented cell lines

The conditional knock-down cell line for *TgSUFC* was generated based on the Tet-Off system using the DHFR-TetO7Sag4 plasmid (75) using a CRISPR-based strategy. Using the DHFR-TetO7Sag4 plasmid as a template, a PCR was performed with the KOD DNA polymerase (Novagen) to amplify the promoter and the resistance gene expression cassette with primers ML4107/ML4108 that also carry 30□bp homology with the 5□ end of the *TgSUFC* gene. A specific single-guide RNA (sgRNA) was generated to introduce a double-stranded break at the 5□ of the *TgSUFC* locus. Primers used to generate the guide were ML4109/ML4110 and the protospacer sequences were introduced in the pU6-Universal plasmid (Addgene ref#52694) (38). The TgSUFC-HA cell line was transfected with the donor sequence and the Cas9/guide RNA-expressing plasmid and transgenic parasites were selected with pyrimethamine and cloned by serial limiting dilution.

The cKD TgSUFC-HA cell line was complemented by the addition of an extra copy of the *TgSUFC* gene put under the dependence of a tubulin promoter at the *uracil phosphoribosyltransferase (UPRT)* locus. *TgSUFC* (1,557 bp) whole cDNA sequence was amplified by reverse transcription (RT)-PCR with primers ML4815/ML4816. They were then cloned downstream of the *tubulin* promoter sequence of the pUPRT-TUB-Ty vector (25) to yield the pUPRT-TgSUFC plasmid. This plasmid was then linearized prior to transfection of the mutant cell line. The recombination efficiency was increased by cotransfecting with the Cas9-expressing pU6-UPRT plasmids generated by integrating (*UPRT*-specific protospacer sequences (with primers ML2087/ML2088 for the 3’ and primers ML3445/ML3446 for the 5’) which were designed to allow a double-strand break at the *UPRT* locus. Transgenic parasites were selected using 5-fluorodeoxyuridine and cloned by serial limiting dilution to yield the cKD TgSUFC-HA comp cell line.

### Anti-TgPDH-E2 antibody production

A polyclonal antibody was raised in rabbit against a peptide (ISLIQAKGLSLISASSSPA) specific of TgPDH-E2 by the Proteogenix company. The peptide was conjugated to Keyhole limpet haemocyanin carrier protein prior to immunization and the whole serum was affinity-purified against the peptide for increased specificity.

### Immunoblot analysis

Protein extracts from 10^7^ freshly egressed tachyzoites were prepared in Laemmli sample buffer, separated by SDS-PAGE and transferred onto nitrocellulose membrane using the BioRad Mini-Transblot system according to the manufacturer’s instructions. Rat monoclonal antibody (clone 3F10, Roche) was used to detect HA-tagged proteins. Other primary antibodies used were mouse monoclonal anti-TY tag (27), rabbit anti-lipoic acid antibody (ab58724, Abcam), mouse anti-SAG1 (76), rabbit anti-CPN60 (77), mouse anti-actin (78), and rabbit anti-farnesyl polyclonal antibody (PA1-12554, Life Technologies).

### Immunofluorescence microscopy

For immunofluorescence assays (IFA), intracellular tachyzoites grown on coverslips containing HFF monolayers, were either fixed for 20 min with 4% (w/v) paraformaldehyde in PBS and permeabilized for 10 min with 0.3% Triton X-100 in PBS or fixed for 5 min in cold methanol (for SAG labelling). Slides/coverslips were subsequently blocked with 0.1% (w/v) BSA in PBS. Primary antibodies used (at 1/1,000, unless specified) were rat anti-HA tag (clone 3F10, Roche), mouse anti-TY tag (27), rabbit anti-CPN60 (77), rabbit anti-IMC3 (79), mouse anti-SAG1 (76), and mouse anti-SAG3 (80). Staining of DNA was performed on fixed cells by incubating them for 5 min in a 1 μg/ml 4,6-diamidino-2-phenylindole (DAPI) solution. All images were acquired at the Montpellier RIO imaging facility from a Zeiss AXIO Imager Z2 epifluorescence microscope driven by the ZEN software v2.3 (Zeiss). Z-stack acquisition and maximal intensity projection was performed to quantify apicoplast loss. Adjustments for brightness and contrast were applied uniformly on the entire image.

### Electron Microscopy

Parasites were pretreated for three days with ATc, and then used to infect HFF monolayers and grown for an extra 24 hours in ATc. They were fixed with 2.5% glutaraldehyde in cacodylate buffer 0.1 M pH7.4. Coverslips were then processed using a Pelco Biowave pro+ (Ted Pella). Briefly, samples were postfixed in 1% OsO_4_ and 2% uranyl acetate, dehydrated in acetonitrile series and embedded in Epon 118 using the following parameters: Glutaraldehyde (150 W ON/OFF/ON 1-min cycles); two buffer washes (40 s 150 W); OsO_4_ (150 W ON/OFF/ON/OFF/ON 1-min cycles); two water washes (40 s 150 W); uranyl acetate (100 W ON/OFF/ON 1-min cycles); dehydration (40 s 150 W); resin infiltration (350 W 3-min cycles). Fixation and infiltration steps were performed under vacuum. Polymerization was performed at 60°C for 48 hr. Ultrathin sections at 70 nM were cut with a Leica UC7 ultramicrotome, counterstained with uranyl acetate and lead citrate and observed in a Jeol 1400+ transmission electron microscope from the MEA Montpellier Electron Microscopy Platform. All chemicals were from Electron Microscopy Sciences, and solvents were from Sigma.

### Plaque assay

Confluent monolayers of HFFs were infected with freshly egressed parasites, which were left to grow for 7□days in the absence or presence of ATc (unless stated). For some experiments, the medium was supplemented with 50 μM palmitic acid (P0500, Sigma-Aldrich), 50 μM myristic acid (70082, Sigma-Aldrich), or 20 μM geranylgeranyol (G3278, Sigma-Aldrich). They were then fixed with 4% v/v paraformaldehyde (PFA) and plaques were revealed by staining with a 0.1% crystal violet solution (V5265, Sigma-Aldrich).

### Gliding assay

10^7^ freshly egressed parasites were resuspended in 300 μl of motility buffer (Ringer’s solution: 155 mM NaCl, 3 mM KCl, 2 mM CaCI_2_, 1 mM MgCl_2_, 3 mM NaH_2_PO_4_, 10 mM HEPES, 10 mM glucose). 100 μl were deposited on poly-L-lysine coated microscope slides (J2800AMNZ, Thermo Scientific), in a well delineated with a hydrophobic pen (PAP Pen, Kisker Biotech). Parasites were left to glide for 15 minutes in an incubator at 37°C, then the suspension was carefully removed and parasites were fixed with 4% (w/v) paraformaldehyde in PBS. Immunostaining was performed with an anti-SAG1 antibody (76) as described above, but without permeabilization. Trail deposition images were acquired with a 63x objective on a Zeiss AXIO Imager Z2 epifluorescence microscope and processed with ImageJ v. 1.53f51, using the NeuronJ plugin as described previously (35).

### Lipidomic analyses

Parasite lipidomic analyses were conducted as previously described (44, 45). Briefly, the parasites were grown for 72 h in +/− ATc conditions within a confluent monolayer of HFF in flasks (175 cm^2^). At each time point, parasites were harvested as intracellular tachyzoites (1 × 10^7^ cell equivalents per replicate) after syringe filtration with 3-μm pore size membrane. These parasites were metabolically quenched by rapid chilling in a dry ice-ethanol slurry bath and then centrifuged down at 4°C. The parasite pellet was washed with ice-cold PBS thrice, before transferring the final pellet to a microcentrifuge tube. Then total lipids were extracted in chloroform/methanol/water (1:3:1, v/v/v) containing PC (C13:0/C13:0), 10 nmol and C21:0 (10 nmol) as internal standards for extraction. Polar and apolar metabolites were separated by phase partitioning by adding chloroform and water to give the ratio of chloroform/methanol/water as 2:1:0.8 (v/v/v). For lipid analysis, the organic phase was dried under N_2_ gas and dissolved in 1-butanol to obtain 1μl butanol/10^7^ parasites.

*Total lipid analysis* - The extracted total lipid sample was then added with 1 nmol pentadecanoic acid (CI5:O) as internal standard as stated before using Trimethylsulfonium hydroxide for total FA content. Resultant FA methyl esters (FAMEs) were analyzed by GC-MS as previously described (43). All FAMEs were identified by comparison of retention time and mass spectra from GC-MS with authentic chemical standards. The concentration of FAMEs was quantified after initial normalization to different internal standards and finally to parasite number.

Stable isotope metabolic labeling experiment.

*Tracking host-derived FAs- (monitoring parasite scavenging capacities)*

*Tracking host-derived fatty acids - (monitoring parasite scavenging capacities)* - Stable isotope metabolic labelling combined to lipidomic analyses have been conducted as previously established and described (45). Briefly, the HFF cells were grown (1 × 10^8^ cell equivalents per replicate) to confluency in the presence of stable U-^13^C-glucose isotope at a final concentration of 800 μM added to a glucose-free DMEM. These ^13^C-pre labelled HFF were then infected with TgNFS2/TgSUFC cKD parasites in the presence of normal-glucose containing DMEM under +/-ATc (0.5 μg/ml). The host HFF and parasites were metabolically quenched separately, and their lipid content was quantified by GC-MS as described above. As described previously, the degree of the incorporation of ^13^C into fatty acids (%carbon incorporation) is determined by the mass isotopomer distribution (MID) of each FAMEs. The total abundance of ^13^C-labelled fatty acids was analyzed initially for HFF to check labelling of the metabolites (described previously). Later, the same was calculated for parasites to confirm direct uptake of ^13^C-labelled fatty acids from the host.

## Data availability

All data are contained within the manuscript. Material described is available upon request from the corresponding author.

## Supporting information

This article contains supporting information.

## Acknowledgments

*We* are grateful to B. Striepen, L. Sheiner, S. Lourido, D. Soldati-Favre, M.J. Gubbels, V. Carruthers, and J.F. Dubremetz for sharing antibodies, strains and plasmids. We wish to thank K. Semenovskaya for technical help with some constructs. We also thank the Montpellier Rio Imaging facility for providing access to their microscopes, as well as the electron microscopy imaging facility of the University of Montpellier.

## Funding and additional information

AC was supported by a fellowship from the Fondation pour la Recherche Médicale (Equipe FRM EQ.20170336725), as well as CYB, CLV and YYB (Equipe FRM EQU202103012700). SB and CYB acknowledge support from the Labex Parafrap (ANR-11-LABX-0024), the Agence Nationale de la Recherche (ANR-21-CE44-0010 to CYB and SB, and ANR-19-CE15-0023 to SB). Funding from the Région Auvergne Rhone-Alpes for the lipidomics analyses platform is also acknowledged (Grant I RICE Project GEMELI). The funders had no role in study design, data collection and analysis, decision to publish, or preparation of the manuscript.

## Abbreviations

ABC: (ATP-binding cassette),
ATc: (anhydrotetracycline),
cKD: (conditional knock-down),
DMAPP: (dimethylallyl diphosphate),
DOXP: (1-deoxy-D-xylulose 5-phosphate),
ER: (endoplasmic reticulum),
ETC: (electron transport chain),
FA: (fatty acid),
FAME: (fatty acid methyl ester),
FASII: (type II fatty acid synthase),
FBS: (fetal bovine serum),
Fd: (ferredoxin),
Fe-S: (iron-sulfur),
GC-MS: (gas chromatographymass spectrometry),
GGOH: (geranylgeraniol),
GPI: (glycosylphosphatidylinositol),
HA: (hemagglutinin),
HFF: (human foreskin fibroblasts),
IFA: (immunofluorescence assay),
IMC: (inner membrane complex),
IPP: (isopentenyl diphosphate),
ISC: (iron-sulfur cluster),
LipA: (lipoic acid synthase A),
LOPIT: (localization of organelle proteins by isotope tagging),
PDH: (pyruvate dehydrogenase),
surface antigen: (SAG),
SUF: (sulfur utilization factor),
TATi: (tetracycline-inducible transactivator),
UPRT: (uracil phosphoribosyltransferase)

## Supporting Information

This article contains supporting information (Figures S1-S6, Table SI)

