## Supplementary information for "Disrupting the plastid-hosted iron-sulfur cluster biogenesis pathway in *Toxoplasma gondii* has pleiotropic effects irreversibly impacting parasite viability"

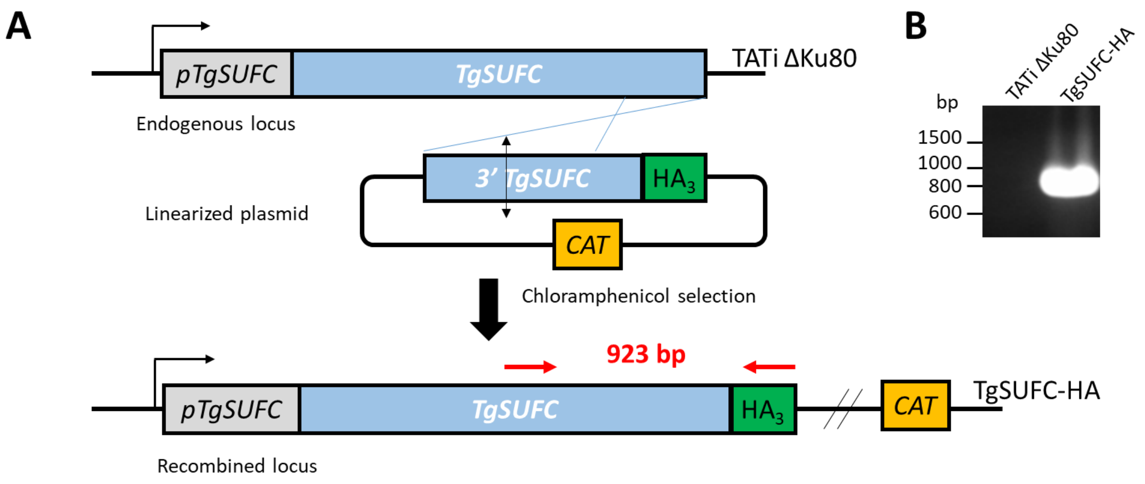


**Figure S1. Generation of an HA-tagged TgSUFC cell line.** A) Schematic representation of the strategy for expressing HA-tagged versions of TgSUFC by homologous recombination at the native *TgSUFC* locus. Chloramphenicol was used to select transgenic parasites based on their expression of the Chloramphenicol acetyltransferase (CAT). B) Diagnostic PCR for verifying correct integration of the construct. The amplified fragments correspond to the red arrows in A), and specific primers used were ML3984/ML1476.

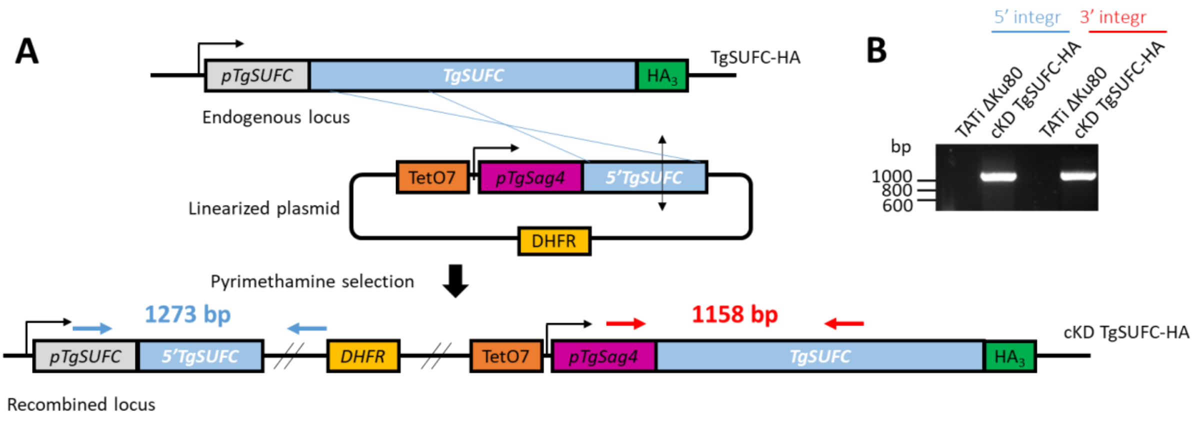


**Figure S2. Generation of a TgSUFC conditional mutant.** A) Schematic representation of the strategy for generating a TgSUFC conditional knock-down cell line by homologous recombination at the native locus. Pyrimethamine was used to select transgenic parasites based on their expression of Dihydrofolate reductase (DHFR). B) Diagnostic PCR for verifying correct integration of the construct. The amplified fragments confirming 5' and 3' integration correspond to the blue and red arrows displayed in A), respectively, and specific primers used were: ML4111/ML687 (5' integration), and ML1041/ML4112 (3' integration).

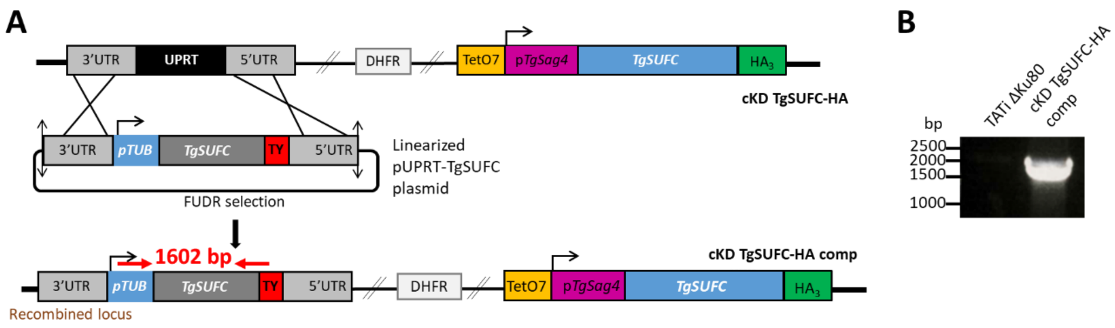


**Figure S3. Generation of a TgSUFC complemented cell line.** A) Schematic representation of the strategy for generating a TgSUFC complemented cell line by integrating an extra copy of the gene of interest by double homologous recombination at the *Uracil Phosphoribosyltransferase* (*UPRT*) locus. Negative selection with 5-fluorodeoxyuridine (FUDR) was used to select transgenic parasites based on their absence of UPRT expression. B) Diagnostic PCR for verifying correct integration of the construct. The amplified fragment confirming integration correspond to the red arrows displayed in A), and specific primers used were ML801/ML4816.

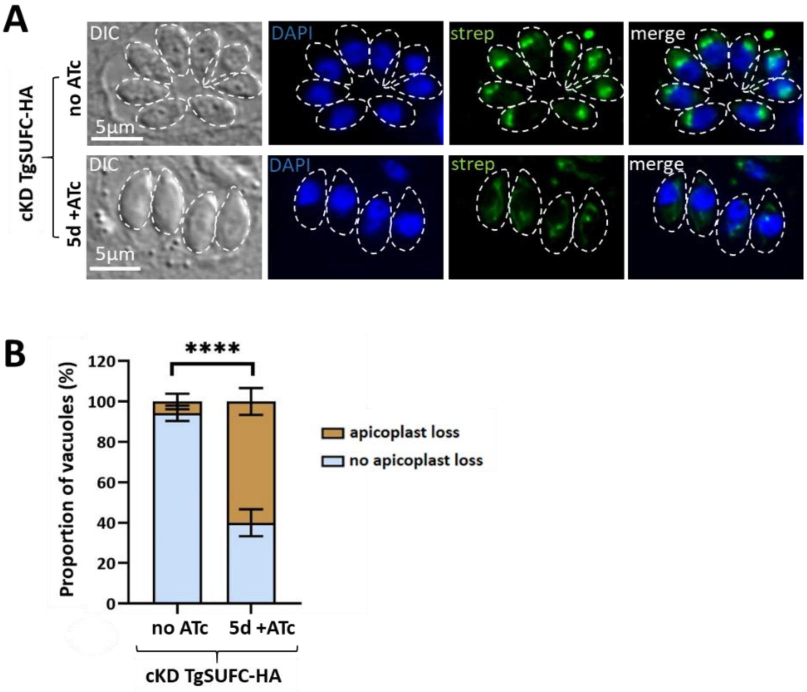


**Figure S4. Apicoplast loss in upon depletion of TgSUFC monitored with streptavidin.** A) Typical streptavidin-Fluorescein Isothiocyanate labelling of the apicoplast (green) vacuoles containing cKD TgSUFC-HA parasites (outlined) grown in the presence or absence of ATc for 5 days; DNA was stained with DAPI (blue); DIC: differential interference contrast. B) Percentage of cKD TgSUFC-HA parasites-containing vacuoles displaying a loss of apicoplast signal when labelled with streptavidin-Fluorescein Isothiocyanate after culture in the presence or absence of ATc for 5 days. Data are mean values from $n = 3$ independent experiments ±SEM. **** $p \leq 0.0001$, Student's *t*-test.

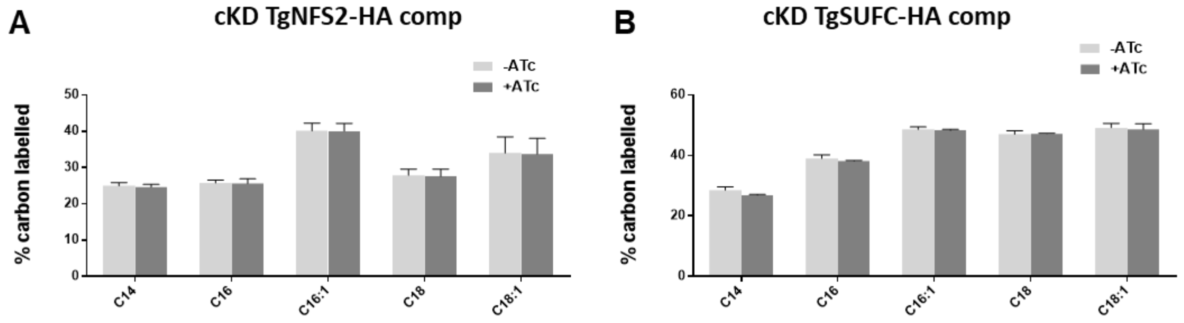


**Figure S5. No major change in host-derived FA uptake in the complemented SUF mutant cell lines.** Host-scavenged lipid flux analyses by stable isotope labelling combined to gas chromatography/mass spectrometry analyses on the complemented TgNFS2-HA (A) and TgSUFC-HA mutant cell lines (B). Data are mean from *n*=3 independent experiments ±SEM.

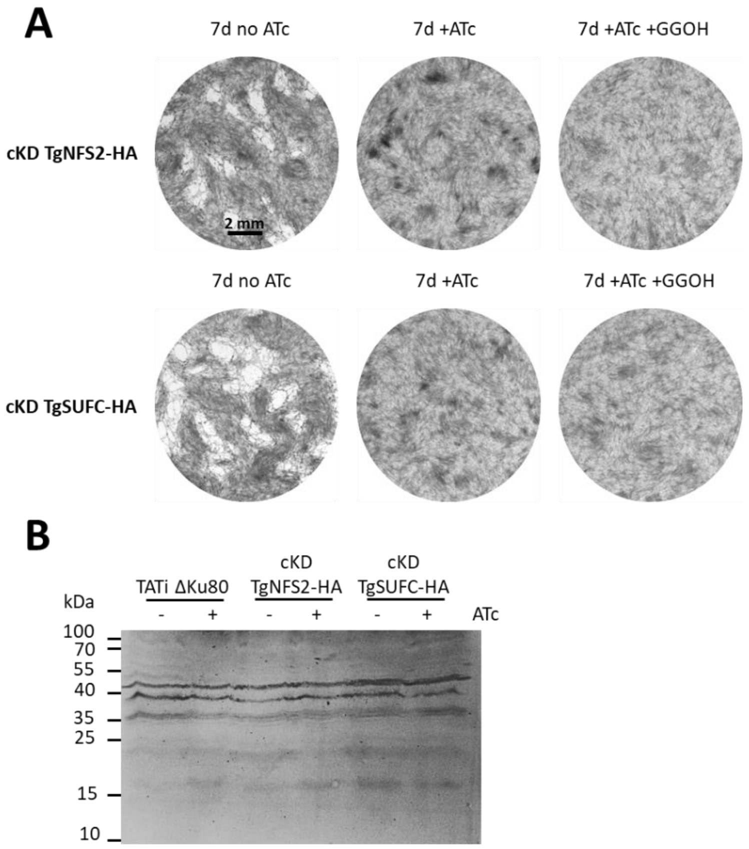


**Figure S6. Depletion of either TgNFS2 or TgSUFC has no strong visible impact on protein farnesylation or prenylation.** A) Plaque assays were performed in the presence or absence of ATc or of 20 µM geranylgeraniol (GGOH). GGOH supplementation did not restore parasite growth in parasites depleted of TgNFS2 or TgSUFC. B) Depletion of TgNFS2 or TgSUFC upon incubation of cKD parasites with ATc for three days has no obvious impact on the global *T. gondii* prenylation profile, as analyzed by mmunoblot probed with an anti-farnesyl antibody. Specific prenylated species are currently unidentified in *T. gondii*.

| Primer name | Primer sequence |
|---|---|
| ML687 | GTTTGAATGCAAGGTTTCGTGCTGTCG |
| ML801 | GAAGACATCCACCAAACGG |
| ML1041 | CGGATCATTTGAAAACATCGTGAGGCTGG |
| ML1476 | CAGCGTAGTCCGGGACGTCGTAC |
| ML2087 | AAGTGCTCCCACGTCCCTCACCAT |
| ML2088 | AAAAATGGTGAGGGACGTGGGAGC |
| ML3445 | AAGTCAGGGCTTCTAAAATGGCGC |
| ML3446 | AAAAGCGCCATTTTAGAAGCCCTG |
| ML3952 | AAGTGTTGAGGTACGATGTGAACAT |
| ML3953 | AAAAATGTTCACATCGTACCTCAAC |
| ML3980 | GCGGCAGACGCAATCTAGACCGCCTGCTCTACCCGTACGACGTCC |
| ML3981 | GGTTTGGCATGAAATTACTGTTACACCGCTCTAGAACTAGTGGATCCCC |
| ML3984 | CGATCTGAGTGCACAGCGAGCC |
| ML4010 | GCCGGATCCTTAGAGCAGGCGGTCTAG |
| ML4107 | TCTGTCTTCCGCGAACCTGAGTTCCGCTTGAAGCTTCGCCAGGCTGTA |
| ML4108 | CTCAACAAGGAACACTGGTAGCGACGCCATAGATCTGGTTGAAGACAGAC |
| ML4109 | AAGTGTGCCACGCCTTCTCATCAAG |
| ML4110 | AAAACTTGATGAGAAGGCGTGGCAC |
| ML4111 | GTCGGACACATCGCACACAGCG |
| ML4112 | TCTGTCTGTGTTTCGTTAGAGCC |
| ML4200 | GCCAAGCTTCATGCCGCTCCTCGAGATCAAAG |
| ML4815 | TTTAGATCTATGGCGTCGCTACCAGTG |
| ML4816 | TTTCCTAGGGAGCAGGCGGTCTAGATTG |
| ML4816 | TTTCCTAGGGAGCAGGCGGTCTAGATTG |

**Table S1. Primers used in this study.**
